# A Mixed-Model Approach for Powerful Testing of Genetic Associations with Cancer Risk Incorporating Tumor Characteristics

**DOI:** 10.1101/446039

**Authors:** Haoyu Zhang, Ni Zhao, Thomas U. Ahearn, William Wheeler, Montserrat García-Closas, Nilanjan Chatterjee

## Abstract

Cancers are routinely classified into subtypes according to various features, including histopathological characteristics and molecular markers. Previous genome-wide association studies have reported heterogeneous associations between loci and cancer subtypes. However, it is not evident what is the optimal modeling strategy for handling correlated tumor features, missing data, and increased degrees-of-freedom in the underlying tests of associations. We propose to test for genetic associations using a mixed-effect two-stage polytomous model score test (MTOP). In the first stage, a standard polytomous model is used to specify all possible sub-types defined by the cross-classification of the tumor characteristics. In the second stage, the subtype-specific case-control odds ratios are specified using a more parsimonious model based on the case-control odds ratio for a baseline subtype, and the case-case parameters associated with tumor markers. Further, to reduce the degrees-of-freedom, we specify case-case parameters for additional exploratory markers using a random-effect model. We use the Expectation-Maximization (EM) algorithm to account for missing data on tumor markers. Through simulations across a range of realistic scenarios and data from the Polish Breast Cancer Study (PBCS), we show MTOP outperforms alternative methods for identifying heterogeneous associations between risk loci and tumor subtypes. The proposed methods have been implemented in a user-friendly and high-speed R statistical package called TOP (https://github.com/andrewhaoyu/TOP).

## I. INTRODUCTION

Genome-wide association studies (GWAS) have identified hundreds of single nucleotide polymorphisms (SNPs) associated with various cancers (MacArthur *and others*, 2016). However, many cancer GWAS have often defined cancer endpoints according to specific anatomic sites, and not according to subtypes of the disease. Many cancers consist of etiologically and clinically heterogeneous subtypes that are defined by multiple correlated tumor characteristics. For instance, breast cancer is routinely classified into subtypes defined by tumor expression of estrogen receptor (ER), progesterone receptor (PR), and human epidermal growth factor receptor 2 (HER2) (Perou *and others*, 2000; Prat *and others*, 2015).

Increasing numbers of epidemiologic studies with tumor specimens are allowing the characterization of cancers at the histological and molecular levels (Cancer Genome Atlas Research, 2014; Network, 2012), providing tremendous opportunities to investigate for potential distinct etiological pathways between cancer subtypes. For example, a breast cancer ER-negative specific GWAS reported 20 SNPs that were more strongly associated with the risk of developing ER-negative than ER-positive disease (Milne *and others*, 2017). Previous studies also suggested traditional breast cancer risk factors, such as age, obesity, and hormone therapy use, were heterogeneously associated with the risk of breast cancer subtypes (Barnard *and others*, 2015).

The most common procedure for testing for associations between risk factors and cancer subtypes is by fitting a standard logistic regression for each subtype versus a control group, then accounting for multiple testing. However, this procedure has several limitations. First, it’s common for cancer cases to have missing tumor marker data, leading to many cancer cases with no subtype definition, and often these cases are dropped from the model. Second, the tumor markers that defined the subtypes are commonly highly correlated with each other. Testing each subtype separately without modeling the correlation limits the power of the model. Finally, as the number of tumor markers increases, the number of cancer subtypes dramatically increases, thus the increased degrees of freedom penalizes the power of the model.

A two-stage polytomous logistic regression was previously proposed to characterize sub-type heterogeneity of a disease according to the underlying disease characteristics (Chatterjee, 2004). The first stage of this method uses a polytomous logistic regression (Dubin and Pasternack, 1986) to model subtype-specific case-control odds ratios. In the second stage, the subtype-specific case-control odds ratios are decomposed into a case-control odds ratio for a reference subtype, a case-case odds ratio for each tumor characteristic, and higher-order interactions between the tumor characteristics. The two-stage model can reduce the degrees of freedom by constraining some or all of the higher-order interactions to be 0. Moreover, the second stage case-case odds ratios can be interpreted as the measures of etiological heterogeneity for tumor characteristics.

Although the two-stage model can improve the power compared to fitting standard logistic regressions for each subtype (Chatterjee, 2004; Zabor and Begg, 2017), the two-stage model does have notable limitations and has not been widely applied to analyze data on multiple tumor characteristics. First, similar to standard logistic regression, the two-stage model can not handle missing tumor characteristics, which is common in epidemiologic studies. Second, the two-stage model estimation algorithm places high demands on computing power and is therefore not readily applicable to large datasets. Finally, although the two-stage model can reduce the multiple testing burdens compared to traditional methods, as the number of tumor characteristics increases, the two-stage model can still have substantial power loss due to the degrees of freedom penalty.

In this paper, we propose a series of computational and statistical innovations to perform computationally scalable and statistically efficient association tests in large cancer GWASs that incorporate tumor characteristic data. Within this two-stage modeling framework, we propose three alternative types of hypotheses for testing genetic associations in the presence of tumor heterogeneity. As the degrees of freedom for the tests can be large in the presence of many tumor characteristics, we propose modeling parameters associated with exploratory tumor characteristics using a random-effect model. We then derive the score tests under the resulting mixed-effect model while taking into account missing data on tumor characteristics using an efficient EM algorithm (Dempster *and others*, 1977). All combined, our work represents a conceptually distinct and practically important extension of earlier methods based on mixed-/fixed-effect models (Lin, 1997; Sun *and others*, 2013; Wu *and others*, 2011; Zhang and Lin, 2003) to the novel setting of modeling genetic associations with multiple tumor characteristics.

The paper is organized as follows. In Section **??**, we describe the proposed three different hypothesis tests, the missing data algorithm, and the score tests. In Section III, we present the simulation results for type I error, power and computation time. In Section IV, the proposed methods are illustrated with applications using data from the Polish Breast Cancer Study (PBCS). In Section V, we discuss the strengths and limitations of the methods and future research directions.

## II. TWO-STAGE POLYTOMOUS LOGISTIC MODEL

The details of the two-stage polytomous logistic model have been described earlier (Chatterjee, 2004). We briefly summarize them for completeness. Suppose a disease can be classified using *K* disease characteristics, and each characteristic *k* can be classified into *M*_*k*_ categories; thus, the disease can be classified into *M* ≡ *M*_1_ × *M*_2_ … × *M*_*K*_ subtypes. For example, breast cancer can be classified into eight subtypes by three tumor characteristics (ER, PR, and HER2), each of which is defined as either positive or negative.

Let *D*_*i*_ denote the disease status of subject *i* in the study such that *D*_*i*_ ∈ {0, 1, 2, …, *M*} and *i* ∈ {1, …, *N*}. *D*_*i*_ = 0 represents a control, and *D*_*i*_ = *m* represents a case with disease subtype *m*. Let *G*_*i*_ be the genotype for subject *i*, and **X**_*i*_ be a *P* × 1 vector of other covariates, where P is the total number of other covariates. In the first stage model, a “saturated” polytomous logistic regression model is constructed as follows:

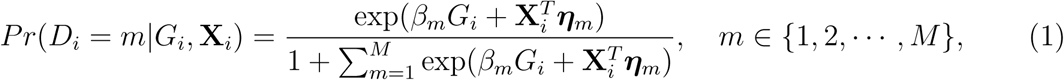

where *β*_*m*_ and ***η***_*m*_ are the regression coefficients for the SNP and other covariates with the *m*th subtype, respectively.

Because each cancer subtype is defined through a unique combination of the *K* tumor characteristics, we can always alternatively index the parameters *β*_*m*_ as 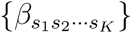, where *s*_*k*_ ∈ {0, 1} for binary tumor characteristics, and 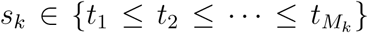 for ordinal tumor characteristics with 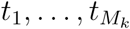 as a set of ordinal scores for *M*_*k*_ different levels. With this new index, the log odds ratios in the first stage can be represented as follows:

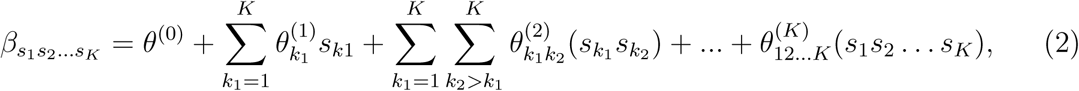

where *θ*^(0)^ represents the case-control log odds ratio for a reference disease subtype, 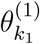 represents the main effect of *k*_1_th tumor characteristic, 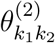 represents the second order interaction between *k*_1_th and *k*_2_th tumor characteristics, and so on. A reference level can be defined for each tumor characteristic, and the reference disease subtype is jointly defined by the combination of the K tumor characteristics.

The reparameterization in 2 provides a way to decompose the first stage parameters to a lower dimension. We can constrain different main effects or interaction effects to be 0 to specify different second stage models. The first stage and second stage parameters can be linked with a matrix form, 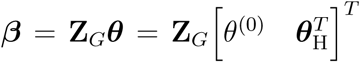, where ***β*** = (*β, β*, …, *β*_*M*_)^*T*^ is a vector of first stage case-control log odds ratios for all the M subtypes, *θ*^(0)^ is the case-control log odds ratio for a reference subtype, and ***θ***_H_ is a vector containing the main effects and interactions effects in the second stage. We will refer to ***θ***_H_ as case-case parameters, and 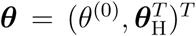 as the vector of second stage parameters. **Z**_*G*_ is the second stage design matrix connecting the first stage and second stage parameters. By constraining different second stage main effects or interaction effects to be 0, we can construct different **Z**_*G*_ to build different two-stage models.

Up to now, we have only described second stage decomposition for the regression coefficients of **G**. The second stage decomposition can also be applied to the other covariates, the details of which are in Supplementary Section 1. We suggest not to perform second stage decomposition on the intercepts parameters of the first stage polytomous model, i.e., the coefficients of intercepts are saturated, because decomposing the intercepts equates to making assumptions on the prevalence of different cancer subtypes, which can potentially lead to bias. Moving forward, we use **Z**_**X**_ to denote the second stage design matrix for the other covariates **X, λ** to denote the second stage parameters for **X**, and **Z** to denote the second stage design matrix for all the covariates.

### A. Hypothesis test under two-stage model

The first stage case-control log odds ratios of subtypes can be decomposed into the second stage case-control log odds ratio of the reference subtype, main effects and interaction effects of tumor characteristics. This decomposition presents multiple options for comprehensively testing for the association between a SNP and cancer subtypes. The first hypothesis test is the global association test, 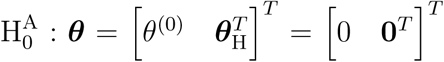 versus 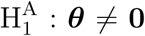, which tests for an overall association between the SNP and the disease. Because ***θ*** = **0** implies ***β*** = **0**, rejecting this null hypothesis means the SNP is associated with at least one of the subtypes. The null hypothesis can be rejected if the SNP is significantly associated with a similar effect size across all subtypes (i.e. ***θ***^(0)^ ≠ 0, ***θ***_H_ = **0**), or if the SNP has heterogeneous effects on different subtypes (***θ***_H_ ≠ **0**).

The second hypothesis test is the global heterogeneity test, 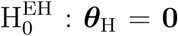 versus 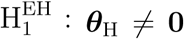. This test simultaneously evaluates the etiologic heterogeneity with respect to a SNP and all the tumor characteristics. Rejecting this null hypothesis indicates that the first stage case-control log odds ratios are significantly different between at least two different subtypes.

Notably, the global heterogeneity test does not identify which tumor characteristic(s) is/are driving the heterogeneity. To identify the tumor characteristic(s) responsible for observed heterogeneity, we propose the individual tumor marker heterogeneity test, 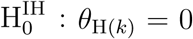 versus 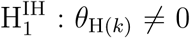, where *θ*_H(*k*)_ is one of the case-case parameters of ***θ***_H_. The case-case parameter (*θ*_H(*k*)_) provides a measurement of etiological heterogeneity according to a specific tumor characteristic (Begg and Zhang, 1994). In the breast cancer example, we can directly test 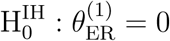 versus 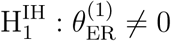. Rejecting the null hypothesis provides evidence that the case-control log odds ratios of ER+ and ER-subtypes are significantly different.

### B. EM algorithm accounting for cases with incomplete tumor characteristics

In the previous sections, all the tumor characteristics were assumed to have no missing data. However, in epidemiological research, it is very common to have missing tumor characteristics. This problem becomes exacerbated as the number of tumor characteristics grows. Restricting to cases with complete tumor characteristics can reduce statistical power and potentially introduce selection bias. To solve this problem, we propose to use the EM algorithm (Dempster *and others*, 1977) to find the maximum likelihood estimate (MLE) of the two-stage model, while incorporating all available information from the study. Let **T**_*io*_ be the observed tumor characteristics of subject *i*, and *Y*_*im*_ = *I*(*D*_*i*_ = *m*) denote whether the *i*th subject is disease subtype *m*. Given **T**_*io*_, the possible subtypes for subject *i*, denoted as 𝒴_*io*_ = {*Y*_*im*_ : *Y*_*im*_ that is consistent with **T**_*io*_}, are within a limited subset of all possible tumor subtypes. We assume that (*Y*_*i*1_, *Y*_*i*2_, …, *Y*_*iM*_, *G*_*i*_, **X**_*i*_) are independently and identically distributed (i.i.d.), and that the tumor characteristics are missing at random (MAR). Let ***δ*** = (***θ***^*T*^, **λ**^*T*^)^*T*^ represent the second stage parameters of both **G** and **X**. Given the notation, the E step of them EM algorithm at the *v*th iteration is

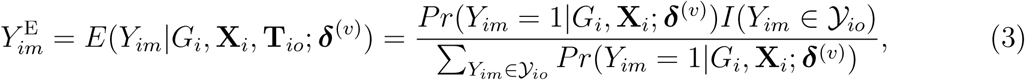

where 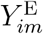 is the probability of the *i*th person to be the *m*th subtype given his observed tumor characteristics (**T**_*io*_), genotype (*G*_*i*_), and other covariates (**X**_*i*_). *I*(*Y*_*im*_ ∈ 𝒴_*io*_) denotes whether the *m*th subtype for the *i*th subject belong to the subsets of possible subtypes given the observed tumor characteristics. The M step at the *v*th iteration is

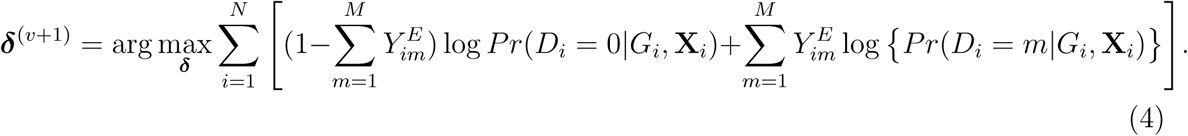

The M step can be solved through a weighted least square iteration. Let **Y**_*m*_ = (*Y*_1*m*_, …, *Y*_*Nm*_)^*T*^, and 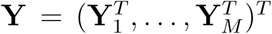. Let **C** = (**G, X**), and **C**_*M*_ = **I**_*M*_ ⊗ **C**. Let **W** = **D** − **AA**^*T*^, **D** = diag(**P**), **P** = *E*(**Y**|**C**; *δ*), and **A** = **D**(**1**_*M*_ ⊗ **I**_*N*_). During the *t*th iteration of the weighted least square, **Y**∗^(*t*)^ = **W**^(*t*)^(**Y**^E^ − **P**^(*t*)^) + **C**_*M*_ **Z*δ***^(*t*)^, where **P**^(*t*)^ and **W**^(*t*)^ are respectively defined as **P** and **W** evaluated at the ***δ***^(*t*)^. The weighted least square update is 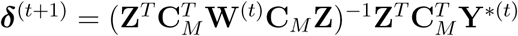. As *t* → ∞, the weighted least square interaction converges to 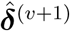, which will be used in next iteration. The EM algorithm will converge to the MLE of the second stage parameters (denoted as 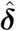), and the observed information matrix **I** is 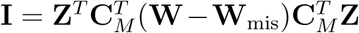, where 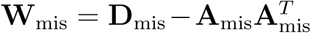, **D**_mis_ = diag(**P**_mis_), **P**_mis_ = *E*(**Y**|**C, T**_*o*_; *δ*), and **A**_mis_ = **D**_mis_(**1**_*M*_ ⊗ **I**_*N*_) (Louis, 1982). More details of the EM algorithm are in Supplementary Section 2.

With the MLE of the second stage parameters of G as 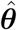, we can construct the the Wald statistics as 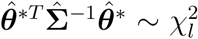 for the global association test, global etiological heterogeneity test, and individual tumor characteristic heterogeneity test using the corresponding second stage parameters and covariance matrix, where the degrees of freedom *l* equal the length of 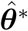.

### C. Fixed-effect two-stage polytomous model score test (FTOP)

Although the hypothesis tests can be implemented through the Wald test, estimating the model parameters for all SNPs in the genome is time-consuming and computationally intensive. In this section, we develop a score test for the global association test assuming the second stage parameters to be fixed. The score test only needs to estimate the second stage parameters of **X** under the null hypothesis once, making it much more computationally efficient than the Wald test. Moreover, the EM algorithm only needs to be implemented once under the null hypothesis. Since we don’t perform any second stage decomposition on the intercept parameters in the first stage polytomous model, the correlations between the tumor characteristics are kept close to the empirical correlations for tumor markers. Most of the imputation power is due to the high correlation between the tumor markers. In the breast cancer example, the correlation between ER and PR is 0.63, between ER and HER2 is −0.16, and between PR and HER2 is −0.17 (Supplementary Table 1). Also, The association of **X** with the tumor markers can improve the power of the EM algorithm. Since a single SNP **G** usually has a small effect, the fact that the effect of individual **G** is not incorporated in the EM algorithm itself doesn’t result in much loss of efficiency.

**TABLE I.** Type one error estimates of MTOP, FTOP with 2.4 × 10^7^ randomly simulated samples. Global test for association and global test for heterogeneity were applied with FTOP and MTOP. Heterogeneity test for a tumor marker was applied with only FTOP. All of the type error rates are divided by the *α* level.

| Interested tests | Total sample size | MTOP |  |  | FTOP |  |  |
| --- | --- | --- | --- | --- | --- | --- | --- |
| | | $\alpha = 10^{-4}$ | $\alpha = 10^{-5}$ | $\alpha = 10^{-6}$ | $\alpha = 10^{-4}$ | $\alpha = 10^{-5}$ | $\alpha = 10^{-6}$ |
| Global association test | 5,000 | .99 | .97 | .88 | .91 | .91 | .67 |
|  | 50,000 | .98 | 1.0 | 1.0 | .99 | 1.0 | .93 |
|  | 100,000 | 1.0 | .94 | 1.0 | 1.0 | 1.0 | 1.0 |
| Global heterogeneity test | 5,000 | 1.0 | .97 | .89 | .92 | .85 | .55 |
|  | 50,000 | 1.0 | 1.0 | 1.0 | 1.0 | 1.0 | 1.0 |
|  | 100,000 | 1.0 | .94 | 1.0 | 1.0 | .98 | .97 |
| Heterogeneity test for<br>a tumor marker | 5,000 |  |  |  | .92 | .93 | .76 |
|  | 50,000 |  |  |  | .98 | .97 | 1.0 |
|  | 100,000 |  |  |  | 1.0 | .97 | 1.0 |

Let **G**_*M*_ = **I**_*M*_ ⊗**G**, and **X**_*M*_ = **I**_*M*_ ⊗**X**. Under the null hypothesis, H_0_ : ***θ*** = **0**, let 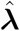 denote the MLE of **λ** under the null hypothesis. The efficient score of ***θ*** is 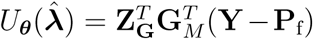, where 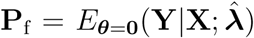. Let 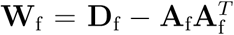, with 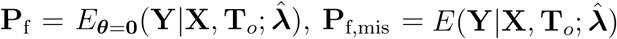, **D**_f_ = diag(**P**_f_ − **P**_f,mis_) and **A**_f_ = **D**_f_(**1**_*M*_ ⊗ **I**_*N*_). The corresponding efficient information matrix of 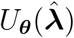 is

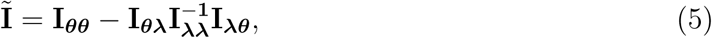

where 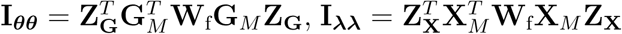, and 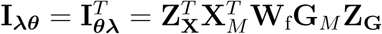.

The score test statistic *Q*_***θ***_ for fixed-effect two stage model is

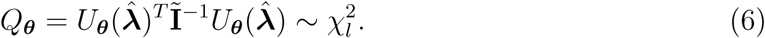

FTOP has the same degrees of freedoms and similar asymptotic power (Yi and Wang, 2011) as the Wald test. In GWAS which needs to perform millions of tests, FTOP can be first used to scan the whole genome with global association test, and then select the potential risk regions. In the selected risk regions, each SNP can be tested for global heterogeneity and individual tumor characteristic heterogeneity using Wald test.

### D. Mixed-effect two-stage polytomous model score test (MTOP)

The two-stage model decreases the degrees of freedom compared to the polytomous logistic regression. However, the power gains in the two-stage model can be reduced as additional tumor characteristics are added into the model. We further propose a mixed-effect two-stage model by modeling some of the second stage case-case parameters as random effects. Let **u** = (*u*_1_, …, *u*_*s*_)^*T*^, where each *u*_*j*_ follows an arbitrary distribution *F* with mean zero and variance *σ*^2^. The mixed-effect second stage model links the first and second stage parameters as follows:

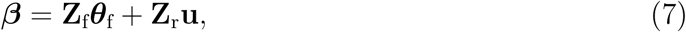

where **Z**_f_ is the second stage design matrix of fixed effect, **Z**_r_ is the second stage design matrix of random effect, and ***θ***_f_ are the fixed-effect second stage parameters. Let 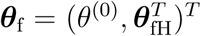, where *θ*^(0)^ is the case-control log odds ratio of the reference subtype, and ***θ***_fH_ are the fixed case-case parameters. The baseline effect *θ*^(0)^ is always kept fixed, since it captures the SNP’s overall effect on all the cancer subtypes.

The fixed-effect parameters ***θ***_fH_ can be used for tumor characters with prior information suggesting that they are a source of heterogeneity, and the random-effect parameters **u** can model tumor characteristics with little or no prior information. In the breast cancer example, the baseline parameter (*θ*^(0)^) and the main effect of ER (*θ*_fH_) can be modeled as fixed effects, since previous evidence indicates ER as a source of breast cancer heterogeneity (García-Closas *and others*, 2013; Milne *and others*, 2017). The main effects of PR and HER2 and other potential interactions effects can be modeled as random effects (**u**). In the mixed effect two-stage model, the global association test is 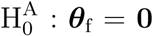, *σ*^2^ = 0 versus 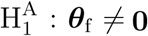 or *σ*^2^ ≠ 0, and the global etiology heterogeneity test is 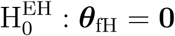, *σ*^2^ = 0 versus 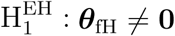 or *σ*^2^ ≠ 0.

To derive the score statistic for the global null 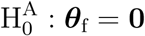, *σ*^2^ = 0, the common approach is to take the partial derivatives of loglikelihood with respective to ***θ***_f_ and *σ*^2^ respectively. However, under the null hypothesis, the score for ***θ***_f_ follows a normal distribution, and for *σ*^2^ follows a mixture of chi-square distribution (Supplementary Section 3). With the correlation between the two scores, getting the joint distribution between the two becomes very complicated. Inspired by methods for the rare variants testing (Sun *and others*, 2013), we propose to modify the derivations of score statistic so that two independent scores can be independent. First for ***θ***_f_, the score test statistic 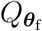 is derived under the global null hypothesis 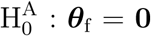, *σ*^2^ = 0 as usual. But for *σ*^2^, the the score statistic 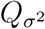 is derived under the null hypothesis H_0_ : *σ*^2^ = 0 without constraining ***θ***_f_. Through this procedure, the two score test statistics (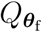 and 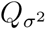) can be proved to be independent (Supplementary Section 4), and the Fisher’s procedure (Koziol and Perlman, 1978) can be used to combine the p-value generated from the two independent tests. Similarly to FTOP, the EM algorithm under the null hypothesis of MTOP can efficiently handle the missing tumor marker problems given the high correlations between the tumor characteristics. However, since MTOP needs to estimate ***θ***_f_ under the null hypothesis H_0_ : *σ*^2^ = 0 for every single SNP, the computation speed for MTOP is slower than FTOP.

The score statistic of the fixed effect ***θ***_f_ under the global null 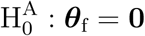, *σ*^2^ = 0 is

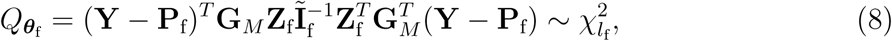

where 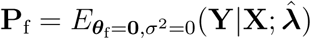. Here 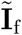 has the same definition as Equation 5, but substitute **Z**_**G**_ with **Z**_f_. Under the null hypothesis, 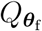 follows a *χ*^2^ distribution with the degrees of freedom *l*_f_ the same as the length of ***θ***_f_.

To explicitly express 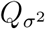, let 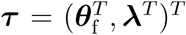 be the second stage fixed effect, and **Z**_***τ***_ is the corresponding second stage design matrix. The variance component score statistic of *σ*^2^ under the null hypothesis H_0_ : *σ*^2^ = 0 without constraining ***θ***_f_ is as follows:

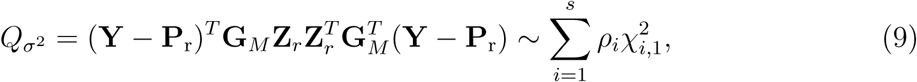

where 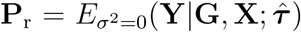, and 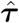 is the MLE under the null hypothesis, H_0_ : *σ*^2^ = 0. Under the null hypothesis, 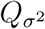 follows a mixture of chi square distribution (Supplementary Section 3), where 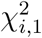 i.i.d. follows 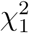. (*ρ*_1_, …, *ρ*_*s*_) are the eigenvalues of 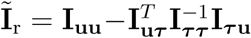, with 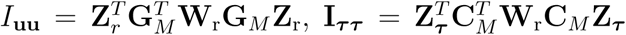 and 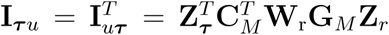, where 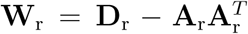, with 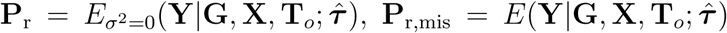, **D**_r_ = diag(**P**_r_ − **P**_r,mis_) and **A**_r_ = **D**_r_(**1**_*M*_ ⊗ **I**_*N*_). The Davies exact method (Davies, 1980) is used here to calculate the p-value of the mixture of chi square distribution.

Let 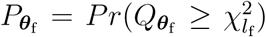 and 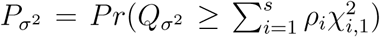 be the p-values of the two independent score statistics. Under the null hypothesis 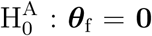, *σ*^2^ = 0, following the Fisher’s procedure, 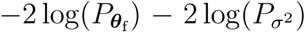 follows 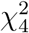; thus, the p-value of mixed effect two-stage model under the null hypothesis is

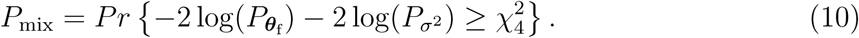

The extension of the score statistics of the global etiology heterogeneity test, 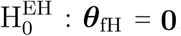, *σ*^2^ = 0, can be computed following a similar procedure as the global association test.

## III. SIMULATION EXPERIMENTS

Large scale simulations across a wide range of practical scenarios were conducted to evaluate the type I error (Section III A), statistical power (Section III B), and computation time (Supplementary Section 5) of the fixed-effect and mixed-effect two-stage models. Data were simulated to mimic the PBCS. We simulated four tumor characteristics: ER (positive vs. negative), PR (positive vs. negative), HER2 (positive vs. negative), and grade (ordinal 1, 2, 3), which collectively defined 2^3^ × 3 = 24 breast cancer subtypes.

In each simulation, genotype data **G** was simulated under the Hardy-Weinberg equilibrium with minor allele frequency (MAF) as 0.25. An additional covariate (**X**) was simulated following a standard normal distribution independent of **G**. We simulated a multinomial outcome with 25 groups, one for the control group, and the other 24 for different cancer subtypes, using the polytomous logistic regression model as follows:

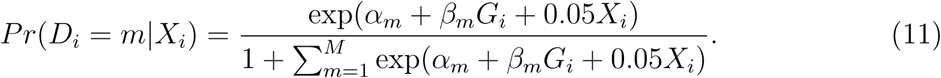

The effect of **X** was set as 0.05 for all subtypes. Using the frequency of the breast cancer subtypes from Breast Cancer Association Consortium (Supplementary Table 2) (Michailidou *and others*, 2017), we computed the corresponding polytomous logistic regression intercept parameters *α*_*m*_. The case-control ratio was set around 1:1, and the proportions of ER+, PR+ and HER2+ were 0.81, 0.68, and 0.17, respectively. The proportions of grade 1, 2, and 3 were 0.20, 0.48, and 0.32. The missing tumor markers were selected randomly with missing rates of 0.17, 0.25, 0.42, and 0.27 for ER, PR, HER2 and grade, respectively. Under this simulation, approximately 70% cases had at least one missing tumor characteristic.

**TABLE II.** Analysis results of previously identified susceptibility loci. For the listed eight loci, MTOP global association test p value decreased more than ten fold compared to the standard logistic regression p value. All of the loci are significant in global heterogeneity test (P *<* 0.05).

| SNP | Chr.<sup>a</sup> | Position | MAF<sup>b</sup> | Global association P | Standard analysis P | Global heterogeneity P |
| --- | --- | --- | --- | --- | --- | --- |
| rs4973768 | 3 | 27,416,013 | .47 | $3.1 \times 10^{-2}$ | $9.5 \times 10^{-1}$ | $9.5 \times 10^{-3}$ |
| rs10816625 | 9 | 110,837,073 | .06 | $5.0 \times 10^{-2}$ | $9.8 \times 10^{-1}$ | $2.2 \times 10^{-2}$ |
| rs7904519 | 10 | 114,773,927 | .46 | $6.5 \times 10^{-2}$ | $8.5 \times 10^{-1}$ | $3.1 \times 10^{-2}$ |
| rs554219 | 11 | 69,331,642 | .13 | $7.3 \times 10^{-11}$ | $1.4 \times 10^{-7}$ | $5.1 \times 10^{-6}$ |
| rs11820646 | 11 | 129,461,171 | .40 | $1.5 \times 10^{-2}$ | $8.6 \times 10^{-1}$ | $4.5 \times 10^{-3}$ |
| rs2236007 | 14 | 37,132,769 | .21 | $2.1 \times 10^{-3}$ | $1.9 \times 10^{-1}$ | $3.5 \times 10^{-3}$ |
| rs1436904 | 18 | 24,570,667 | .40 | $7.2 \times 10^{-4}$ | $6.6 \times 10^{-2}$ | $9.7 \times 10^{-4}$ |
| rs1436904 | 22 | 29,121,087 | .01 | $9.8 \times 10^{-3}$ | $1.6 \times 10^{-1}$ | $2.3 \times 10^{-2}$ |

<sup>a</sup>Chr. chromosome. <sup>b</sup> MAF, minor allele frequency.

### A. Type I error

We evaluated the type I error of the global association test, global heterogeneity test, and individual tumor marker heterogeneity test under the global null hypothesis. The data were generated by setting *β*_*m*_ = 0 in Equation 11, where none of the subtypes was associated with genotypes. The total sample size *n* was set to be 5,000, 50,000 and 100,000. We conducted 2.4 × 10^7^ simulations to evaluate the type I error at *α* = 1.0 × 10^−4^, 1.0 × ×10^−5^, and 1.0 × 10^−6^ level.

Both MTOP and FTOP were applied with an additive two-stage model by constraining all the interaction terms as 0 in Equation 2. The subtype-specific case-control log ORs were specified into the case-control log OR of a baseline disease subtype (ER-, PR-, HER2-, grade 1) and the main effects associated with the four tumor markers. Furthermore, the MTOP assumed the baseline and ER case-case parameter as fixed effects and the other case-case parameters as random effects. The global association test and global heterogeneity test were implemented using both MTOP and FTOP, but the individual tumor characteristic heterogeneity test could only be implemented with FTOP. For MTOP and FTOP, we removed all the subtypes with fewer than 10 cases to avoid potential nonconvergence of the model.

Table I presents the estimated type I errors under the global null hypothesis. Both MTOP and FTOP correctly control the type I error, especially for the larger sample sizes. FTOP is conservative with 5,000 subjects, especially for *α* = 1.0 × 10^−6^, however, the method is still valid. The well-controlled type I error also shows that removing rare subtypes doesn’t bias the estimate, as further demonstrated by additional simulations that are presented in Supplementary Section 6. In the later sections, we generally used the additive second stage structure for both MTOP and FTOP unless otherwise specified.

### B. Statistical power

We assessed the statistical power of the proposed methods using various simulation settings with sample sizes as 25,000, 50,000, and 100,000. For each setting, we performed 2 × 10^5^ simulations to evaluate the power at *α* = 5.0 × 10^−8^ level.

#### 1. Global association test

The data were simulated with three different scenarios: I. no heterogeneity between tumor markers, II. heterogeneity according to one tumor marker, and III. heterogeneity according to multiple tumor markers. The disease subtypes were generated through Equation 11. Under scenario I, we set *β*_*m*_ as 0.08 for all the subtypes. For scenarios II and III, *β*_*m*_ was simulated following the additive two-stage model. Under scenarios II, datasets were simulated with only ER heterogeneity by setting the case-case parameter for ER as 0.08, and all the other as 0. For scenario III, we simulated a scenario with heterogeneity according to all 4 tumor markers by setting the baseline effect to be 0, the ER case-case parameter to be 0.08, and all the other case-case parameters following a normal distribution with mean 0 and variance 4.0 × 10^−4^. Under this scenario, all tumor characteristics contributed to the subtype-specific heterogeneity. Moreover, to evaluate different methods under a larger number of tumor characteristics, additional simulations were conducted by adding two additional binary tumor characteristics to the previous four tumor characteristic setting. This defined 2^5^ × 3 = 96 cancer subtypes. The two additional tumor characteristics were randomly selected to be missing with 5% missing rate. Under this setting, around 77% of the cases have at least one tumor characteristic missing. We compared the statistical power to detect the overall association using FTOP, MTOP, standard logistic regression, FTOP with only complete data, and polytomous logistic regression. For MTOP, FTOP and polytomous model, we removed all the subtypes with fewer than 10 cases to avoid potential nonconvergence of the model.

Overall, MTOP had robust power under all scenarios (Figure 1). Standard logistic regression had the highest power when there was no subtype-specific heterogeneity (Scenario I), but suffered from substantial power loss when heterogeneity existed between subtypes. MTOP, followed by FTOP, consistently demonstrated the highest power among the five methods when subtype-specific heterogeneity existed (scenarios II and III). The power gain of MTOP over FTOP ranged from 2% to 49%. The power gain was small when there were four tumor characteristics because the difference in the degrees of freedom between MTOP and FTOP was small. However, with six tumor markers, the power gain of MTOP was more apparent owing to the larger difference in the degrees of freedom between the models. FTOP was the least efficient in scenarios with no or little heterogeneity, such as scenarios I and II, but with increasing heterogeneity, such-as scenario III, the power of MTOP and FTOP were more similar.

**FIG. 1.**
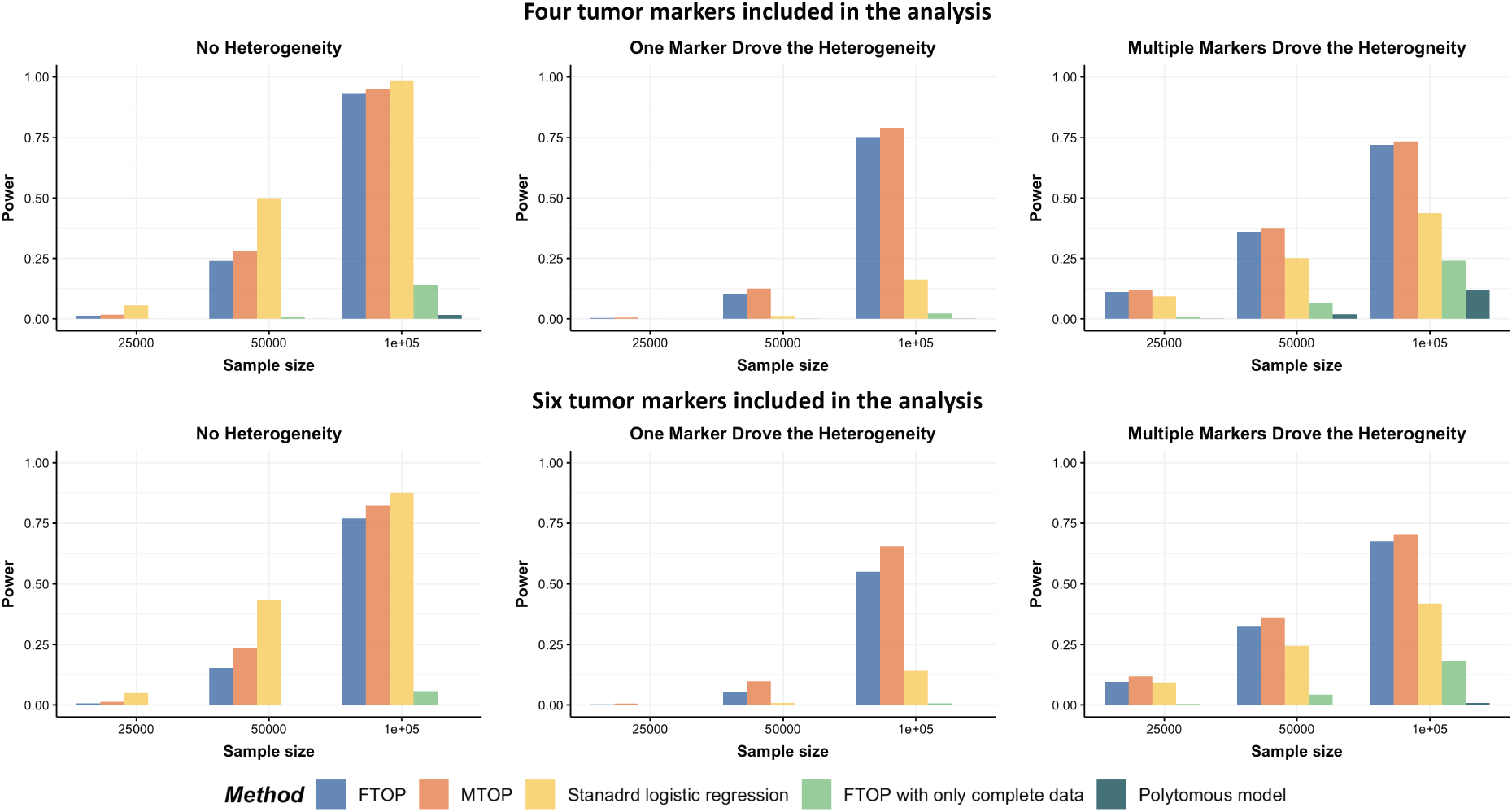
Power comparison among MTOP, FTOP, standard logistic regression, two-stage model with only complete data and polytomous model with 2 × 10^5^ random samples. For the three figures in the first row, four tumor markers were included in the analysis. Three binary tumor marker and one ordinal tumor marker defined 24 cancer subtypes. Around 70% cases would be incomplete. For the three figures in the second row, two extra binary tumor markers were included in the analysis. The six tumor markers defined 96 subtypes. Around 77% cases would be incomplete. The power was estimated by controlling the type one error *α <* 5.0 × 10^−8^.

The simulation study also showed that the incorporation of cases with missing tumor characteristics significantly increased the power of the methods (Figure 1). Under the four tumor markers setting with around 70% incomplete cases, the power gain of FTOP incorporating the missing data algorithm was at least 200% compared to FTOP with only complete data. As expected, under the six tumor markers setting, which resulted in more missing tumor marker data, the power of FTOP with the missing data algorithm was once again significantly higher than FTOP with only complete data. MTOP was the most powerful method when heterogeneity across cancer subtypes was present. Additional power simulations with 5,000 subjects are described in Supplementary Section 7.

The previous simulations mainly focused on the two-stage model with additive effects. Additional simulations were also implemented with pairwise interactions in the model. We simulated data with *β*_*m*_ following a second stage model that included main effects and pairwise interactions as shown in Equation 2 with the case-case parameter for ER 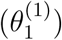 as 0.08, the pairwise interaction effect between ER and HER2 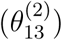 as 0.04, and all the other parameters as 0. Four methods were evaluated including FTOP with/without pairwise interactions and MTOP with/without pairwise interactions (baseline and ER fixed). FTOP without interaction terms still had high power (Figure 2). However, FTOP with pairwise interaction structure had limited power because of the incorporation of the interaction terms as fixed effects. On the other hand, MTOP with/without pairwise interactions maintained a high power even when there were underlying interaction effects.

**FIG. 2.**
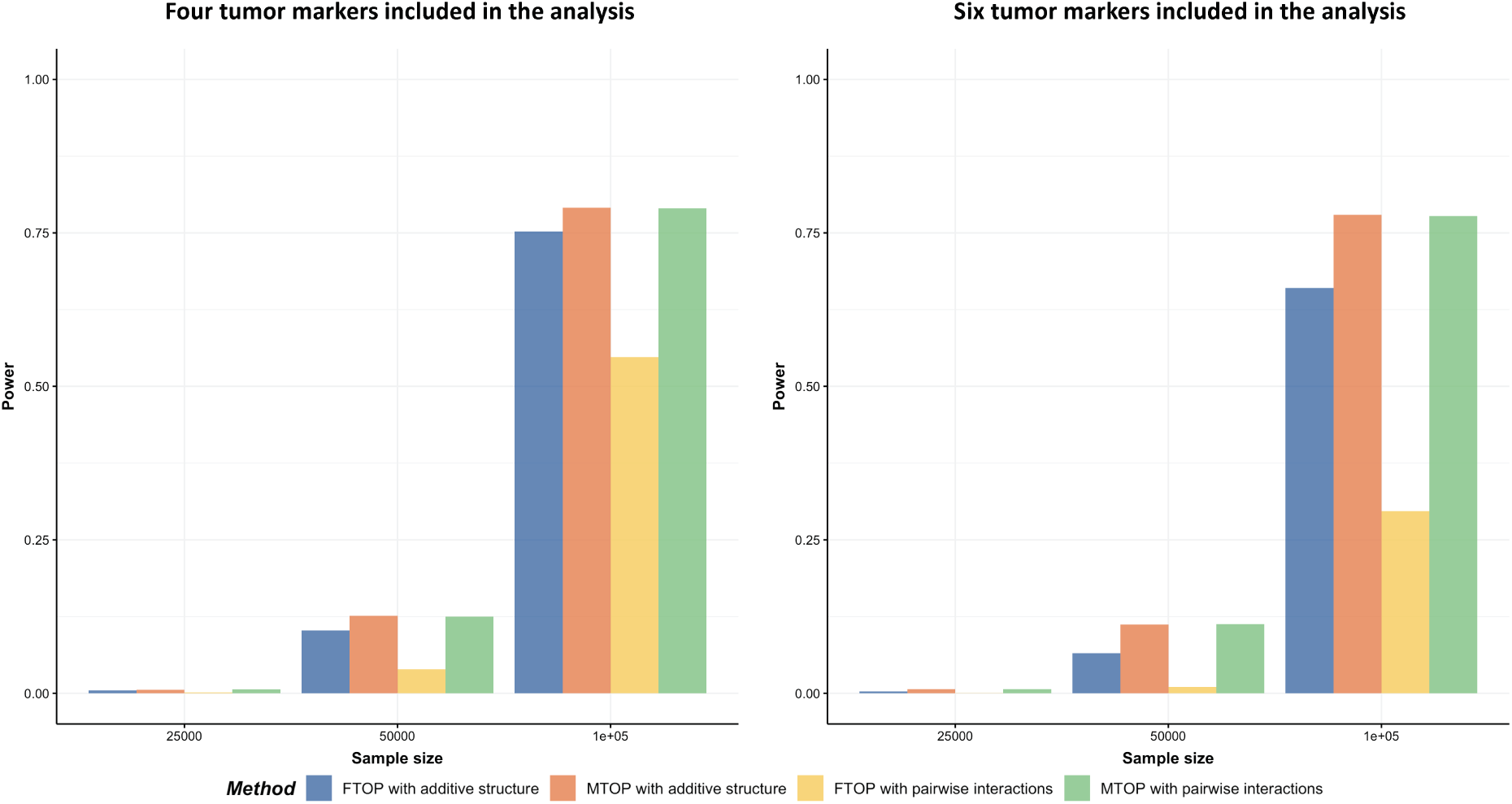
Power comparison of global association test with pairwise interactions. Four methods were evaluated, including FTOP with additive structure, MTOP with additive structure (ER fixed), FTOP with pairwise interactions and MTOP with pairwise interactions (ER fixed). For the three figures in the first row, four tumor markers were included in the analysis. Three binary tumor marker and one ordinal tumor marker defined 24 cancer subtypes. Around 70% cases were incomplete. For the three figures in the second row, two extra binary tumor markers were included in the analysis. The six tumor markers defined 96 subtypes. Around 77% cases were incomplete. The total sample size was 25,000, 50,000 and 100,000. We generated 2 × 10^5^ random replicates. The power was estimated by controlling the type one error *α <* 5.0 × 10^−8^.

**FIG. 3.**
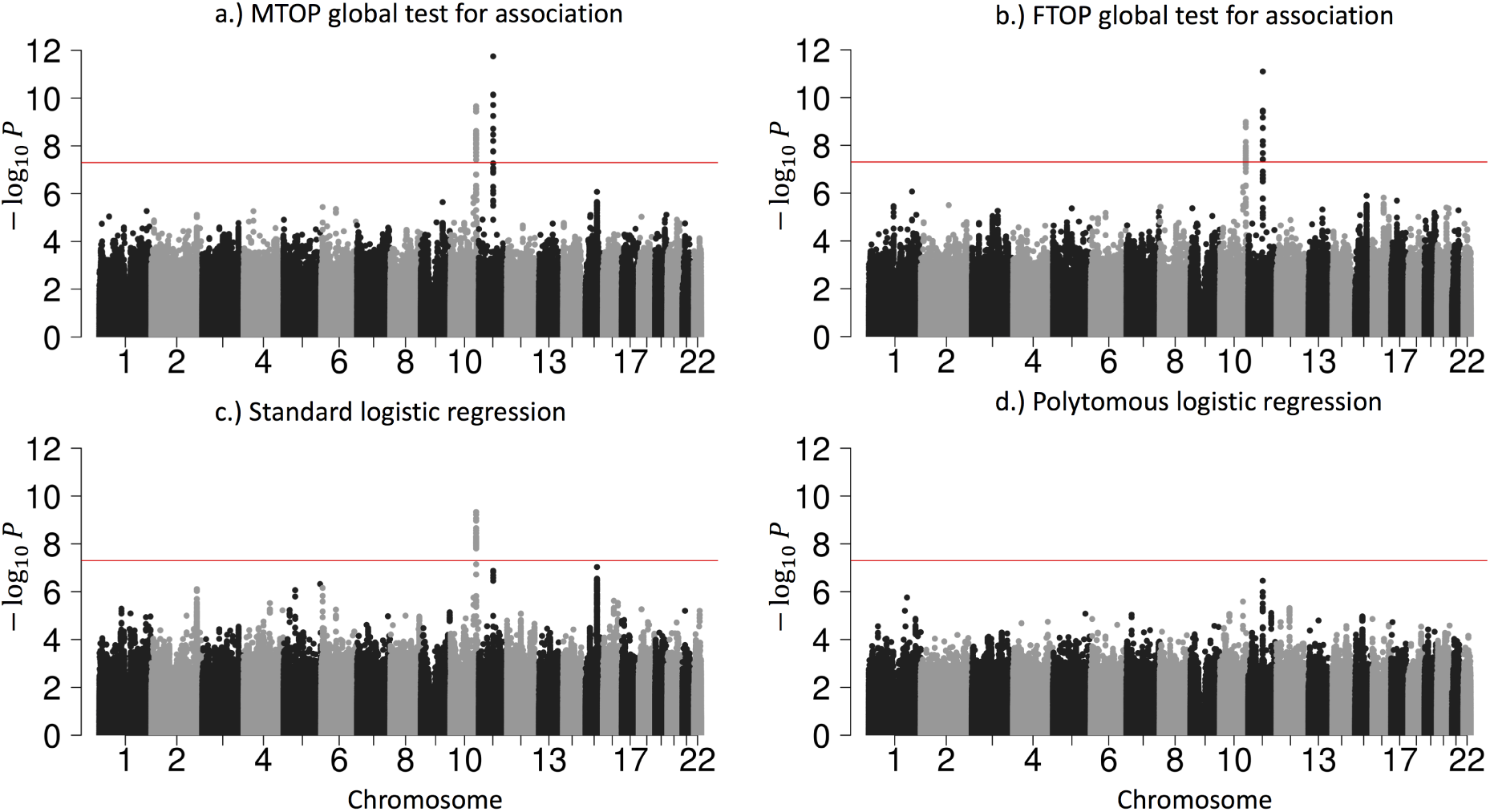
Manhattan plot of genome-wide association analysis with PBCS using four different methods. PBCS have 2,078 invasive breast cancer and 2,219 controls. In total, 7,017,694 SNPs on 22 auto chromosomes with MAF more than 5% were included in the analysis. ER, PR, HER2 and grade were used to define breast cancer subtypes.

#### 2. Global heterogeneity test

Supplementary Figure 3 shows the simulation results for global heterogeneity tests under similar simulation settings as global association tests. MTOP had the highest power when there were heterogeneous associations across the subtypes.

#### 3. Individual tumor marker heterogeneity test

We further evaluated the power of the individual tumor marker heterogeneity test. The data were generated with four tumor characteristics with the ER case-case parameter 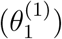 as 0.08, and all other parameters as 0. ER was randomly selected to be missing with a rate of 0.17, 0.30 and 0.50. We compared two different methods, FTOP with all four tumor characteristics and the polytomous model. The polytomous model was set up to test each marker at a time. In the polytomous model, we removed cases with missing data only on the relevant tumor marker to avoid penalizing the power of the model by removing cases that were missing tumor marker data on the other tumor markers. FTOP with all four tumor characteristics had smaller power compared to the polytomous model in testing the effect of ER (Supplementary Figure 4). Since FTOP included all four tumor characteristics, and the tumor markers were highly correlated, the variability of underlying parameters was larger. However, the type I errors of the polytomous model in testing PR, HER2 and grade were inflated under this case (Supplementary Figure 5). Under this simulation, these three markers had no effect. On the other hand, FTOP controlled the type I error of all the tests.

Overall, for the global test for association and the global test for heterogeneity, when there was no heterogeneity, the standard logistic regression was the most powerful method. However, in the presence of subtype heterogeneity, MTOP was the most powerful method, and MTOP had stable power even with a large number of pairwise interactions terms included.

## IV. APPLICATION TO THE POLISH BREAST CANCER STUDY (PBCS)

We applied our proposed methods to the PBCS, a population-based breast cancer case-control study conducted in Poland between 2000 and 2003 (García-Closas *and others*, 2006). The study consisted of 2,078 cases of histologically or cytologically confirmed invasive breast cancer and 2,219 women without a history of breast cancer at enrollment. Information on ER, PR, and grade were available from pathology records (García-Closas *and others*, 2006), and information on HER2 was available from immunohistochemical staining of tissue microarray blocks (Yang *and others*, 2007). We used genome-wide genotyping data to compare MTOP, FTOP, standard logistic regression, and polytomous logistic regression to detect SNPs associated with breast cancer risk.

Supplementary Table 4 presents the sample size of the tumor characteristics. The four tumor characteristics defined 24 mutually exclusive breast cancer subtypes. Subtypes with less than 10 cases were excluded, leaving 17 subtypes in the analysis. Both MTOP and FTOP used the additive second stage design. Besides, we modeled the baseline and ER case-case parameters as fixed effects in MTOP, and all other effects as random effects. We put ER as a fixed effect because of the previously reported heterogeneity in genetic association by ER (García-Closas *and others*, 2013; Milne *and others*, 2017). Genotype imputation was done using IMPUTE2 based on 1000 Genomes Project as reference (Michailidou *and others*, 2017; Milne *and others*, 2017). In total, 7,017,694 common variants on 22 auto chromosomes with MAF ≥ 5% were included in the analysis. In all the models, we adjusted for age and the first four genetic principal components to account for population stratification.

As Figure 3 shows, MTOP, FTOP and standard logistic regression all identified a known susceptibility variant in the FGFR2 locus on chromosome 10 (Michailidou *and others*, 2017), with the most significant SNP being rs11200014 (P *<* 5.0 × 10^−8^). Further, both MTOP and FTOP identified a second known susceptibility locus on chromosome 11 (CCND1) (Michailidou *and others*, 2017), with the most significant SNP in both models being rs78540526 (P *<* 5.0×10^−8^). The individual heterogeneity test of this SNP showed evidence for heterogeneity by ER (P=0.011) and grade (P=0.024). Notably, the CCND1 locus was not genome-wide significant in standard logistic regression or polytomous models. The type I error of the four methods was well-controlled (Supplementary Figure 6).

Additional sensitivity analysis of MTOP was implemented by specifying baseline, ER and grade as fixed effects, and PR and HER2 as random effects (Supplementary Figure 7). The results for MTOP with grade as fixed vs. random effect were similar. We also implemented MTOP and FTOP incorporating pairwise interactions in the second stage model (Supplementary Figure 8-9). With pairwise interactions, both MTOP and FTOP detected FGFR2 and CCND1 with the genome-wide significant threshold. However, the P-value of FTOP with pairwise interactions was less significant compared to FTOP without these interaction terms (for rs11200014, P = 4.3 × 10^−8^ vs. P = 1.0 × 10^−9^; for rs78540526, P= 2.7 × 10^−10^ vs. P = 8.1 × 10^−12^). The P-value of MTOP with pairwise interactions was also less significant compared to MTOP without interaction terms (for rs11200014, P= 1.0 × 10^−9^ vs. P = 2.2 × 10^−10^; for rs78540526, P = 1.7 × 10^−11^ vs. P = 1.8 × 10^−12^). In both scenarios with pairwise interactions parameter included, however, the power loss was smaller.

Next, we compared the ability of MTOP and standard logistic regressions to detect 178 previously identified breast cancer susceptibility loci (Michailidou *and others*, 2017). For eight of the 178 loci, the MTOP global association test p-value was more than ten fold lower compared to the standard logistic regression p-value (Table II). In the MTOP model, these eight loci all had significant global heterogeneity tests (P *<* 0.05). Confirming these results, in a previous analysis applying MTOP to 106,571 breast cancer cases and 95,762 controls, these eight loci were reported to have significant global heterogeneity (Ahearn *and others*, 2019).

## V. DISCUSSION

We present a series of novel methods for performing genetic association testing for cancer outcomes accounting for potential heterogeneity across subtypes. These methods efficiently account for multiple testing, correlations between markers, and missing tumor data. Under the model framework, we develop two computationally efficient score tests, FTOP and MTOP, which model the underlying heterogeneity parameters in terms of fixed effects or mixed effects, respectively. We demonstrate these methods have greater statistical power in the presence of subtype heterogeneity than either standard or polytomous logistic regression analysis.

Several methods have been proposed to study the etiological heterogeneity of cancer subtypes (Chatterjee, 2004; Rosner *and others*, 2013; Wang *and others*, 2015). A recent review showed the well-controlled type I error and good statistical power of the two-stage model (Zabor and Begg, 2017). However, previous two-stage models haven’t accounted for missing tumor markers, which is a common problem in epidemiological studies. We show that by incorporating the EM algorithm into the two-stage model we can take advantage of all available information and substantially increase the statistical power (Figure 1). Moreover, the newly proposed mixed effect model can mitigate the degrees of freedom penalty caused by analyzing many tumor characteristics. In a recent large breast cancer GWAS analysis with 106,571 cases and 95,762 controls, the newly developed methods MTOP and FTOP have identified 16 novel loci (Zhang *and others*, 2019).

Incorporating missing tumor characteristics based on the proposed EM algorithm requires the assumption of MAR, i.e. the mechanism of missing of the individual tumor characteristics can depend only on other observed tumor characteristics and covariates, but not on the unobserved missing value themselves. For the analysis of tumor heterogeneity, information on aggressive types of tumors may be systematically missing. If the missing tumor characteristics are important determinants of aggressiveness, then the underlying assumption is violated. In general, dealing with non-ignorable missing data is a complex problem and certain sensitivity analyses can be performed to explore the degree of bias (Little and Rubin, 2019). In the context of genetic association testing, non-ignorable missingness can lead to inflated type I error only if the missingness mechanism itself is related to the genetic variant. Further research is merited to explore the complex effects of non-ignorable missingness in type I error and power of the proposed tests.

The computation time of MTOP is greater than FTOP (Supplementary Section 5). To construct the score tests in FTOP, the coefficients of covariates need to be estimated once under the null hypothesis, while in MTOP they need to be estimated for every SNP. The computational complexity of FTOP is O(*NM* ^2^*P* ^2^), with *P* as the number of other covariates **X**. For MTOP, the computational complexity is O(*NM* ^2^*P* ^2^lk), where l and k are respectively the numbers of iteration required for weighted least square and EM algorithm to converge.

Currently, we only implement the linear kernel in MTOP, but other common kernels that capture the similarity between tumor characteristics can be used in the future. If there is prior knowledge about the overlapping genetic architecture across different tumor subtypes, this will help to choose the kernel function, and improve the power of the methods.

The proposed methods have been implemented in a user-friendly and high-speed R statistical package called TOP (https://github.com/andrewhaoyu/TOP), which includes all the core functions implemented in C code.

## VI. SUPPLEMENTARY MATERIALS

In Supplementary Section 1, the two-stage model is generalized to multivariates. In Supplementary Section 2-3, the details of the EM algorithm and the variance component score statistic are respectively presented. In Supplementary Section 4, 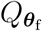 and 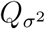 are proved to be independent. In Supplementary Section 5, computation time simulations are presented. In Supplementary Section 6, the simulations to evaluate the bias of the estimates are shown. In Supplementary Section 7, simulations with 5,000 subjects are presented. Supplementary Table 1 shows the correlations of ER, PR, HER2, and grade. Supplementary Table 2 presents the frequencies of the joint distribution of ER, PR, HER2, and grade. Supplementary Table 3 shows the simulation results to evaluate bias. Supplementary Table 4 presents the sample size of tumor characteristics in PBCS. Supplementary Figure 1 shows the computation time simulations results. Supplementary Figure 2 presents the power analysis of the global association test with 5,000 subjects. Supplementary Figure 3 presents global heterogeneity test simulation results. Supplementary 4-5 respectively present the power and type I error simulations results of individual tumor marker heterogeneity test. Supplementary Figure 6 is the QQ plot of GWAS with PBCS. Supplementary Figure 7 shows the GWAS with PBCS using MTOP with ER and grade as fixed effects. Supplementary Figure 8-9 respectively present the GWAS with PBCS using MTOP/FTOP with pairwise interactions.

## ACKNOWLEDGEMENTS

This work was supported by funds from the NCI Intramural Research Program, Bloomberg Distinguished Professorship endowment, and NHGRI (1R01 HG010480-01). The simulation experiments and data analysis were implemented using the high performance computation Biowulf cluster at National Institutes of Health, USA.

