## Supplementary material for "A Mixed-Model Approach for Powerful Testing of Genetic Associations with Cancer Risk Incorporating Tumor Characteristics": supplmentary_material

February 5, 2020

#### 1 Generalize the two-stage model to multivariate

We define $\mathbf{I}_k$ as a $k \times k$ identity matrix, and $\mathbf{1}_q$ as a $q \times 1$ unit vector. Let $\mathbf{G} = (G_1, \ldots, G_N)^T$, and $\mathbf{X} = (\mathbf{X}_1, \ldots, \mathbf{X}_N)^T$. Let $\eta_{mp}$ be the regression coefficient of the $m$th subtype with $p$th covariate. Let $\boldsymbol{\eta}_p^{(c)} = (\eta_{1p}, \ldots, \eta_{Mp})^T$, where $\boldsymbol{\eta}_p^{(c)}$ represents $M$ subtypes first stage parameters of the $p$th covariate. Here we grouped the first stage parameters by covariates, and this notation could be convenient for introducing different second stage design matrices for multiple covariates. $\mathbf{Z}_p$ is the second stage design matrix of $p$th other covariate linking the first stage regression coefficients $\boldsymbol{\eta}_p^{(c)}$ and corresponding second stage parameters $\boldsymbol{\lambda}_p$ for $p$th other covariate, where $\boldsymbol{\eta}_p^{(c)} = \mathbf{Z}_p \boldsymbol{\lambda}_p$. Let $\boldsymbol{\lambda} = (\boldsymbol{\lambda}_1^T, \ldots, \boldsymbol{\lambda}_P^T)^T$ and $\boldsymbol{\eta}^{(c)} = \{\boldsymbol{\eta}_1^{(c)T}, \ldots, \boldsymbol{\eta}_P^{(c)T}\}^T$ , where $\boldsymbol{\lambda}$ and $\boldsymbol{\eta}^{(c)}$are the second stage and first stage parameters for all the P other covariates. Let $\mathbf{Z}_{\mathbf{X}}^{(c)} = \oplus_{p=1}^{P} \mathbf{Z}_p$, where $\oplus_{p=1}^{P} \mathbf{Z}_p$ is a block diagonal matrix with $\mathbf{Z}_1$ to $\mathbf{Z}_P$ as the diagonal block.

For the convenience of estimation procedure, we will use $\boldsymbol{\eta}_m = (\eta_{m1}, \ldots, \eta_{mP})^T$ to represent the $P$ covariates first stage parameters of $m$th subtype. Let $\boldsymbol{\eta} = (\boldsymbol{\eta}_1^T, \ldots, \boldsymbol{\eta}_M^T)^T$, where $\boldsymbol{\eta}$ represent the first stage parameters for the P other covariates grouped by subtypes, and $\boldsymbol{\eta}$ is a reordering of $\boldsymbol{\eta}^{(c)}$. Let $\mathbf{Z}_{\mathbf{X}} = \{\mathbf{Z}_{\mathbf{X}}^{(c)}\}_{\pi}$, where $\pi$ is reordering the row of the design matrix to group the first-stage parameters by subtypes, thus $\boldsymbol{\eta} = \mathbf{Z}_{\mathbf{X}} \boldsymbol{\lambda}$. We don't perform any second stage decomposition on the regression coefficients of intercepts, since making assumption on the prevalence of different cancer subtypes could potentially yield bias, which means $\mathbf{Z}_1 = \mathbf{I}_M$. The second stage parameters are always grouped by covariates.

To combine the notations of the coefficients of genotype $\mathbf{G}$ and other covariates $\mathbf{X}$, we define $\mathbf{d}_m = (\beta_m, \boldsymbol{\eta}_m^T)^T$, where $\mathbf{d}_m$ represent the first stage parameters of both $\mathbf{G}$ and $\mathbf{X}$ for the $m$th subtype, and let $\mathbf{d} = (\mathbf{d}_1^T, \ldots, \mathbf{d}_M^T)^T$ and $\boldsymbol{\delta} = (\boldsymbol{\theta}^T, \boldsymbol{\lambda}^T)^T$, where $\boldsymbol{\delta}$ represent the second stage parameters of both $\mathbf{G}$ and $\mathbf{X}$. Let $\mathbf{Z} = \{\mathbf{Z}_G \oplus \mathbf{Z}_{\mathbf{X}}^{(c)}\}_{\pi}$, then we have $\mathbf{d} = \mathbf{Z}\boldsymbol{\delta}$, where the second stage design matrix $\mathbf{Z}$ links the first stage regression coefficients to the second stage parameters for all of the covariates.

### 2 Two-stage model EM algorithm derivation

Let $\mathbf{Z} = [\mathbf{Z}_1^{*T}, \ldots, \mathbf{Z}_M^{*T}]^T$, where we split the second stage design matrix by rows into $M$ different matrices with $\mathbf{Z}_m^*$ as the $1 + (m-1) \times P$ to $m \times P$ rows of $\mathbf{Z}$. Since $\mathbf{d} = (\mathbf{d}_1^T, \ldots, \mathbf{d}_M^T)^T$ and $\mathbf{d} = \mathbf{Z}\boldsymbol{\delta}$ (demonstrated in Supplementary Section 1), then $\mathbf{d}_m = \mathbf{Z}_m^*\boldsymbol{\delta}$ and $\mathbf{Z}_m^*$ links the first stage parameters of the $m$th subtypes $\mathbf{d}_m$ to the second stage parameters $\boldsymbol{\delta}$. To avoid notation abuse, we will use $\mathbf{Z}_m$ to represent $\mathbf{Z}_m^*$ through the Supplementary sections.

Let $Y_{im} = I(D_i = m)$ denote whether the $i$th subject has subtype $m$ and $\mathbf{T}_{io}$ be the observed tumor characteristics status of the $i$th subject. Given the observed tumor characteristics, the possible subtypes for subject $i$ will be a limited subset of all possible tumor subtypes, which can be denoted as $\mathcal{Y}_{io} = \{Y_{im} : Y_{im} \text{that is consistent with } \mathbf{T}_{io}\}$. Given $\mathbf{C} = (\mathbf{G}, \mathbf{X})$, let $\mathbf{C} = (\mathbf{C}_1^T, \ldots, \mathbf{C}_N^T)^T$, where $\mathbf{C}_i$ is a $(P+1) \times 1$ covariates vector for the $i$th subject. We assume that $(Y_{i1}, Y_{i2}, \ldots, Y_{iM}, \mathbf{C}_i^T)$ are all observed and independently and identically distributed (i.i.d.) distributed. Then the complete data log likelihood $\log L$ is

$$\log L = \sum_{i=1}^{N} \left[ \sum_{j=1}^{M} Y_{ij} \mathbf{C}_i^T \mathbf{Z}_j \boldsymbol{\delta} - \log\left\{1 + \sum_{m=1}^{M} \exp(\mathbf{C}_i^T \mathbf{Z}_m \boldsymbol{\delta})\right\} \right]. \tag{1}$$

In the $v$th iteration of the EM algorithm E step, we take the expectation of the latent variables $(Y_{i1}, Y_{i2}, \ldots, Y_{iM})$ given the observation tumor characteristics $\mathbf{T}_{io}$:

**E step:**

$$Y_{im}^{\mathrm{E}} = E(Y_{im}|\mathbf{C}_i, \mathbf{T}_{io}; \boldsymbol{\delta}^{(v)}) = \frac{Pr(Y_{im}|\mathbf{C}_i; \boldsymbol{\delta}^{(v)}) I(Y_{im} \in \mathcal{Y}_{io})}{\sum_{Y_{im} \in \mathcal{Y}_{io}} Pr(Y_{im} = 1|\mathbf{C}_i; \boldsymbol{\delta}^{(v)})},$$

where $Y_{im}^{\mathrm{E}}$ is the probability of the $i$th person to be $m$th subtype given his observed tumor characteristics, genotype and other covariates. $I(Y_{im} \in \mathcal{Y}_{io})$ denote whether the $m$th subtype for the $i$th subject belong to the subsets of possible subtypes given the observed tumor characteristics. We defined the complete data log likelihood with expectation to the observed tumor characteristics as $A(\boldsymbol{\delta}|\boldsymbol{\delta}^{(v)}) = E_{Y|\mathbf{T}_o, \boldsymbol{\delta}^{(v)}}(\log L)$. In the **M step**, we get the updates of $\boldsymbol{\delta}$ by maximizing $A(\boldsymbol{\delta}|\boldsymbol{\delta}^{(v)})$ , $\boldsymbol{\delta}^{(v+1)} = \arg\max_{\boldsymbol{\delta}} A(\boldsymbol{\delta}|\boldsymbol{\delta}^{(v)})$, where

$$A(\boldsymbol{\delta}|\boldsymbol{\delta}^{(v)}) = \sum_{i=1}^{N} \left[ \sum_{j=1}^{M} Y_{ij}^{\mathrm{E}} \mathbf{C}_i^T \mathbf{Z}_j - \log\{1 + \sum_{m=1}^{M} \exp(\mathbf{C}_i^T \mathbf{Z}_m \boldsymbol{\delta})\} \right]. \tag{2}$$

The **M step** could be solved with weighted least square interaction algorithm. Let $\mathbf{Y}_m^{\mathrm{E}} = (Y_{1m}^{\mathrm{E}}, \dots, Y_{Nm}^{\mathrm{E}})^T$, and $\mathbf{Y}^{\mathrm{E}} = \{(\mathbf{Y}_1^{\mathrm{E}})^T, \dots, (\mathbf{Y}_M^{\mathrm{E}})^T\}^T$. By taking derivatives of $A(\boldsymbol{\delta}|\boldsymbol{\delta}^{(v)})$, we have

$$\frac{\partial A(\boldsymbol{\delta}|\boldsymbol{\delta}^{(v)})}{\partial \boldsymbol{\delta}} = \sum_{i=1}^{N} \left\{ \sum_{j=1}^{M} \mathbf{Z}_j^T \mathbf{C}_i Y_{ij}^{\mathrm{E}} - \frac{\sum_{k=1}^{M} \mathbf{Z}_k^T \mathbf{C}_i \exp(\mathbf{C}_i^T \mathbf{Z}_k \boldsymbol{\delta})}{1 + \sum_{m=1}^{M} \exp(\mathbf{C}_i^T \mathbf{Z}_m \boldsymbol{\delta})} \right\}, \tag{3}$$

and

$$\begin{aligned} -\frac{\partial A(\boldsymbol{\delta}|\boldsymbol{\delta}^{(v)})}{\partial \boldsymbol{\delta} \partial \boldsymbol{\delta}^T} = \sum_{i=1}^{N} \Bigg[ & \frac{\sum_{k=1}^{M} \mathbf{Z}_k^T \mathbf{C}_i \exp(\mathbf{C}_i^T \mathbf{Z}_k \boldsymbol{\delta}) \mathbf{C}_i^T \mathbf{Z}_k}{1 + \sum_{m=1}^{M} \exp(\mathbf{C}_i^T \mathbf{Z}_m \boldsymbol{\delta})} \\ & - \frac{\{\sum_{k=1}^{M} \mathbf{Z}_k^T \mathbf{C}_i \exp(\mathbf{C}_i^T \mathbf{Z}_k \boldsymbol{\delta})\}}{1 + \sum_{m=1}^{M} \exp(\mathbf{C}_i^T \mathbf{Z}_m \boldsymbol{\delta})} * \frac{\{\sum_{l=1}^{M} \exp(\mathbf{C}_i^T \mathbf{Z}_l \boldsymbol{\delta}) \mathbf{C}_i^T \mathbf{Z}_l\}}{1 + \sum_{m=1}^{M} \exp(\mathbf{C}_i^T \mathbf{Z}_m \boldsymbol{\delta})} \Bigg]. \end{aligned} \tag{4}$$

By writing the Equation 3 and 4 into matrix form, we have :

$$\frac{\partial A(\boldsymbol{\delta}|\boldsymbol{\delta}^{(v)})}{\partial \boldsymbol{\delta}} = \mathbf{Z}^T \mathbf{C}_M^T (\mathbf{Y}^{\mathrm{E}} - \mathbf{P}), \tag{5}$$

and

$$-\frac{\partial A(\boldsymbol{\delta}|\boldsymbol{\delta}^{(v)})}{\partial \boldsymbol{\delta} \partial \boldsymbol{\delta}^T} = \mathbf{Z}^T \mathbf{C}_M^T \mathbf{W} \mathbf{C}_M \mathbf{Z}, \tag{6}$$

, where $\mathbf{C}_M = \mathbf{I}_M \otimes \mathbf{C}$, and the weighted matrix $\mathbf{W} = \mathbf{D} - \mathbf{A}\mathbf{A}^T$, with $\mathbf{D} = \mathrm{diag}(\mathbf{P} - \mathbf{P}_{\mathrm{mis}})$, $\mathbf{P} = E(\mathbf{Y}|\mathbf{C}; \hat{\delta})$, $\mathbf{P}_{\mathrm{mis}} = E(\mathbf{Y}|\mathbf{C}, \mathbf{T}_o; \hat{\delta})$, and $\mathbf{A} = \mathbf{D}(\mathbf{1}_M \otimes \mathbf{I}_N)$. During the $t$th iteration of the weighted Least square, let $\mathbf{Y}^{*(t)} = \mathbf{W}^{(l)}(\mathbf{Y}^{\mathrm{E}} - \mathbf{P}^{(t)}) + \mathbf{C}_M \mathbf{Z} \boldsymbol{\delta}^{(t)}$, where $\mathbf{P}^{(t)}$ and $\mathbf{W}^{(l)}$ uses the same definition as $\mathbf{P}$ and $\mathbf{W}$ evaluated at the $\boldsymbol{\delta}^{(t)}$. The weighted least square updates would be $\boldsymbol{\delta}^{(t+1)} = (\mathbf{Z}^T \mathbf{C}_M^T \mathbf{W}^{(t)} \mathbf{C}_M \mathbf{Z})^{-1} \mathbf{Z}^T \mathbf{C}_M^T \mathbf{Y}^{*(t)}$ . As $t \to \infty$, the weighted least square interaction converged to $\hat{\boldsymbol{\delta}}^{(c)}$.

### 3 Variance component score statistics $Q_{\sigma^2}$ derivation

Given the mixed effect two-stage model $\boldsymbol{\beta} = \mathbf{Z}_{\mathrm{f}} \boldsymbol{\theta}_{\mathrm{f}} + \mathbf{Z}_{\mathrm{r}} \mathbf{u}$, where we decompose the first stage regression parameters of genotype into fixed effect $\boldsymbol{\theta}_f$ and random effect $\mathbf{u}$. $\mathbf{Z}_{\mathrm{f}}$ and $\mathbf{Z}_{\mathrm{r}}$ are the corresponding design matrix for $\boldsymbol{\theta}_{\mathrm{f}}$ and $\mathbf{u}$. Let $\boldsymbol{\tau} = (\boldsymbol{\theta}_{\mathrm{f}}^T, \boldsymbol{\lambda}^T)^T$ and $\mathbf{Z}_{\boldsymbol{\tau}} = \{\mathbf{Z}_{\mathrm{f}} \oplus (\oplus_{p=1}^{P} \mathbf{Z}^{(p)})\}_{\pi}$, where $\boldsymbol{\tau}$ represents the vector of second stage fixed effects and $\mathbf{Z}_{\boldsymbol{\tau}}$ is the corresponding second stage design matrix. Then let the design matrix $\mathbf{Z} = [\mathbf{Z}_{\boldsymbol{\tau}}, \mathbf{Z}_{\mathrm{r}}]$, where $\mathbf{Z}$ is the second stage design matrix for all of the covariates. Following the designation of $\mathbf{Z}_m$ in Appendix A, we have $\mathbf{Z}_m = [\mathbf{Z}_{m\boldsymbol{\tau}}, \mathbf{Z}_{m\mathrm{r}}]$. Given the assumption that $\mathbf{u}$ follows an arbitrary distribution $F$ with mean 0 and variance $\sigma^2$. The likelihood of the mixed effect two-stage model $L(\boldsymbol{\tau}, \sigma^2)$ would be,

$$L(\boldsymbol{\tau}, \sigma^2) = \int \exp\{l(\boldsymbol{\tau}, \mathbf{u})\} dF(\mathbf{u}; \sigma^2), \tag{7}$$

where

$$l(\boldsymbol{\tau}, \mathbf{u}) = \sum_{i=1}^{N} \Bigg( \sum_{j=1}^{M} Y_{ij} \mathbf{C}_i^T (\mathbf{Z}_{m\boldsymbol{\tau}} \boldsymbol{\tau} + Z_{m\mathrm{r}} \mathbf{u}) - \log \Big[ 1 + \sum_{m=1}^{M} \exp \big\{ \mathbf{C}_i^T (\mathbf{Z}_{m\boldsymbol{\tau}} \boldsymbol{\tau} + Z_{m\mathbf{r}} \mathbf{u}) \big\} \Bigg). \tag{8}$$

By taking a two-term Taylor expansion,

$$\exp\{l(\boldsymbol{\tau}, \mathbf{u})\} = \exp l(\boldsymbol{\tau}, \mathbf{0}) \Big[ 1 + \frac{\partial l(\boldsymbol{\tau}, \mathbf{0})}{\mathbf{u}} \mathbf{u} + \frac{1}{2} \mathbf{u}^T \big\{ \frac{\partial l(\boldsymbol{\tau}, \mathbf{0})}{\partial \mathbf{u}} \frac{\partial l(\boldsymbol{\tau}, \mathbf{0})}{\partial \mathbf{u}^T} + \frac{\partial^2 l(\boldsymbol{\tau}, \mathbf{0})}{\partial \mathbf{u} \mathbf{u}^T} \big\} \mathbf{u} + \epsilon \Big], \tag{9}$$

where the error $\epsilon$ has the third and higher order terms of $\mathbf{u}$. By writing the likelihood in Equation 7 as $L(\boldsymbol{\tau}, \sigma^2) = E[\exp\{l(\boldsymbol{\tau}, \mathbf{u})\}]$ and then take the log, then log-likelihood of the mixed effect two-stage model $\log L(\boldsymbol{\tau}, \sigma^2)$ would be,

$$\log L(\boldsymbol{\tau}, \sigma^2) = l(\boldsymbol{\tau}, \mathbf{0}) + \frac{1}{2} tr \big\{ \frac{\partial l(\boldsymbol{\tau}, \mathbf{0})}{\partial \mathbf{u}} \frac{\partial l(\boldsymbol{\tau}, \mathbf{0})}{\partial \mathbf{u}^T} + \frac{\partial^2 l(\boldsymbol{\tau}, \mathbf{0})}{\partial \mathbf{u} \mathbf{u}^T} \big\} \sigma^2 + o(\sigma^2). \tag{10}$$

Under the null hypothesis $\mathrm{H}_0 : \sigma^2 = 0$, $\hat{\boldsymbol{\tau}}$ is the MLE of $\boldsymbol{\tau}$. Following similar derivation as Equation 3 and 4, we have $U_{\mathbf{u}}(\hat{\boldsymbol{\tau}}) = \frac{\partial l(\hat{\boldsymbol{\tau}}, \mathbf{0})}{\partial \mathbf{u}} = \mathbf{Z}_\mathrm{r}^T \mathbf{G}_M^T (\mathbf{Y} - \mathbf{P}_r)$ and $-\frac{\partial^2 l(\boldsymbol{\tau}, \mathbf{0})}{\partial \mathbf{u} \mathbf{u}^T} = \tilde{\mathbf{I}}_\mathrm{r}$, where$\mathbf{P}_\mathrm{r} = E(\mathbf{Y}|\mathbf{X}; \hat{\boldsymbol{\tau}})$ and $\tilde{\mathbf{I}}_\mathrm{r} = \mathbf{I}_{\mathbf{u}\mathbf{u}} - \mathbf{I}_{\mathbf{u}\boldsymbol{\tau}}^T \mathbf{I}_{\boldsymbol{\tau}\boldsymbol{\tau}}^{-1} \mathbf{I}_{\boldsymbol{\tau}\mathbf{u}}$, with $I_{\mathbf{u}\mathbf{u}} = \mathbf{Z}_r^T \mathbf{G}_M^T \mathbf{W}_\mathrm{r} \mathbf{G}_M \mathbf{Z}_\mathrm{r}$, $\mathbf{I}_{\boldsymbol{\tau}\boldsymbol{\tau}} = \mathbf{Z}_{\boldsymbol{\tau}}^T \mathbf{C}_M^T \mathbf{W}_\mathrm{r} \mathbf{C}_M \mathbf{Z}_{\boldsymbol{\tau}}$ and $\mathbf{I}_{\boldsymbol{\tau}u} = \mathbf{I}_{u\boldsymbol{\tau}}^T = \mathbf{Z}_{\boldsymbol{\tau}}^T \mathbf{C}_M^T \mathbf{W}_\mathrm{r} \mathbf{G}_M \mathbf{Z}_r$. The weighted matrix $\mathbf{W}_\mathrm{r}$ has the same definition as the one used for Equation 6 , but evaluated under the null hypothesis $\mathrm{H}_0 : \sigma^2 = 0$. By taking derivatives of $\log L(\boldsymbol{\tau}, \sigma^2)$ with respect to $\sigma^2$, the efficient score $U_{\sigma^2}(\hat{\boldsymbol{\tau}})$ would be,

$$U_{\sigma^2}(\hat{\boldsymbol{\tau}}) = \frac{1}{2} \big\{ (\mathbf{Y} - \mathbf{P}_\mathrm{r})^T \mathbf{G}_M \mathbf{Z}_\mathrm{r} \mathbf{Z}_\mathrm{r}^T \mathbf{G}_M^T (\mathbf{Y} - \mathbf{P}_\mathrm{r}) - tr(\tilde{\mathbf{I}}_\mathrm{r}) \big\}. \tag{11}$$

By dropping the second term $tr(\tilde{\mathbf{I}}_\mathrm{r})$, we only use the first term as the variance component score statistics,

$$Q_{\sigma^2} = (\mathbf{Y} - \mathbf{P}_\mathrm{r})^T \mathbf{G}_M \mathbf{Z}_\mathrm{r} \mathbf{Z}_\mathrm{r}^T \mathbf{G}_M^T (\mathbf{Y} - \mathbf{P}_\mathrm{r}). \tag{12}$$

$Q_{\sigma^2}$ follows a mixture of chi-square distributions $\sum_{i=1}^{s} \rho_i \chi_{i,1}^2$, where $\chi_{i,1}^2$ i.i.d. follows $\chi_1^2$ and $(\rho_1, \ldots, \rho_s)$ are the eigenvalues of $\tilde{\mathbf{I}}_\mathrm{r}$. Proof is as following, Since the the efficient score of $\mathbf{u}$ under the null hypothesis $\mathrm{H}_0 : \sigma^2 = 0$ is $U_{\mathbf{u}}(\hat{\boldsymbol{\tau}}) = \mathbf{Z}_\mathrm{r}^T \mathbf{G}_M^T (\mathbf{Y} - \mathbf{P}_r)$ , then $U_{\mathbf{u}}(\hat{\boldsymbol{\tau}}) \xrightarrow{d} N(\mathbf{0}, \tilde{I}_\mathrm{r})$. Let the eigen decomposition of $\tilde{\mathbf{I}}_\mathrm{r}$ to be $A^T \Gamma A$, where $\Gamma$ is a diagonal matrix whose entries are the eigenvalues of $\tilde{\mathbf{I}}_\mathrm{r}$, and $A$ is an orthogonal matrix whose columns are eigenvectors of $\tilde{\mathbf{I}}_\mathrm{r}$. $Q_{\sigma^2}$ can be transformed as

$$Q_{\sigma^2} = U_{\mathbf{u}}(\hat{\boldsymbol{\tau}})^T U_{\mathbf{u}}(\hat{\boldsymbol{\tau}}) = U_{\mathbf{u}}(\hat{\boldsymbol{\tau}})^T \tilde{\mathbf{I}}_{\mathrm{r}}^{-\frac{1}{2}} A^T \Gamma A \tilde{\mathbf{I}}_{\mathrm{r}}^{-\frac{1}{2}} U_{\mathbf{u}}(\hat{\boldsymbol{\tau}}), \tag{13}$$

where $\tilde{\mathbf{I}}_{\mathrm{r}}^{-\frac{1}{2}} = A\Gamma^{-\frac{1}{2}}A^T$ and $\Gamma^{-\frac{1}{2}}$ is a diagonal matrix with entries as $(\frac{1}{\sqrt{\rho_1}}, \ldots, \frac{1}{\sqrt{\rho_s}})$. Since$A\tilde{\mathbf{I}}_{\mathrm{r}}^{-\frac{1}{2}} U_{\mathbf{u}}(\hat{\boldsymbol{\tau}}) \xrightarrow{d} N(\mathbf{0}, \mathbf{I}_s)$, where $\mathbf{I}_s$ is an $s \times s$ identity matrix, then $Q_{\sigma^2}$ follows a mixture of chi-square distributions $\sum_{i=1}^{s} \rho_i \chi^2_{i,1}$.

### 4 Proof of independence between $Q_{\boldsymbol{\theta}_{\mathrm{f}}}$ and $Q_{\sigma^2}$

Let $\mathbf{P} = E_{\boldsymbol{\theta}_{\mathrm{f}}=\mathbf{0},\sigma^2=0}(\mathbf{Y}|\mathbf{X};\boldsymbol{\lambda})$ be the mean of $\mathbf{Y}$ under the global null hypothesis $\mathrm{H}_0 : \boldsymbol{\theta}_f = \mathbf{0}, \sigma^2 = 0$. Let $U_{\boldsymbol{\theta}_{\mathrm{f}}}(\hat{\boldsymbol{\lambda}}) = \mathbf{Z}_{\mathrm{f}}^T \mathbf{G}_M(\mathbf{Y} - \mathbf{P}_{\mathrm{f}})$ be the efficient score of $\boldsymbol{\theta}_{\mathrm{f}}$, where $\mathbf{P}_{\mathrm{f}} = E_{\boldsymbol{\theta}_{\mathrm{f}}=\mathbf{0},\sigma^2=0}(\mathbf{Y}|\mathbf{X};\hat{\boldsymbol{\lambda}})$ . Then we have $Q_{\boldsymbol{\theta}_{\mathrm{f}}} = U_{\boldsymbol{\theta}_{\mathrm{f}}}(\hat{\boldsymbol{\lambda}})^T \tilde{\mathbf{I}}_{\mathrm{f}}^{-1} U_{\boldsymbol{\theta}_{\mathrm{f}}}(\hat{\boldsymbol{\lambda}})$ and $Q_{\sigma^2} = U_u(\hat{\boldsymbol{\tau}})^T U_u(\hat{\boldsymbol{\tau}})$, where $\tilde{\mathbf{I}} = \mathbf{I}_{\boldsymbol{\theta}\boldsymbol{\theta}} - \mathbf{I}_{\boldsymbol{\theta}\boldsymbol{\lambda}}^T \mathbf{I}_{\boldsymbol{\lambda}\boldsymbol{\lambda}} \mathbf{I}_{\boldsymbol{\lambda}\boldsymbol{\theta}}$, with $\mathbf{I}_{\boldsymbol{\theta}\boldsymbol{\theta}} = \mathbf{Z}_{\mathbf{G}}^T \mathbf{G}_M^T \mathbf{W}_{\mathrm{f}} \mathbf{G}_M \mathbf{Z}_{\mathbf{G}}$, $\mathbf{I}_{\boldsymbol{\lambda}\boldsymbol{\lambda}} = \mathbf{Z}_{\mathbf{X}}^T \mathbf{X}_M^T \mathbf{W}_{\mathrm{f}} \mathbf{X}_M \mathbf{Z}_{\mathbf{X}}$, and $\mathbf{I}_{\boldsymbol{\lambda}\boldsymbol{\theta}} = \mathbf{I}_{\boldsymbol{\lambda}\boldsymbol{\theta}}^T = \mathbf{Z}_{\mathbf{X}}^T \mathbf{X}_M^T \mathbf{W}_{\mathrm{f}} \mathbf{G}_M \mathbf{Z}_{\mathbf{G}}$. The weighted matrix $\mathbf{W}_{\mathrm{f}}$ has the same definition as in Equation 6, but evaluated under the null hypothesis $\mathrm{H}_0 : \boldsymbol{\theta} = \mathbf{0}$. To prove the independence between $Q_{\boldsymbol{\theta}_{\mathrm{f}}}$ and $Q_{\sigma^2}$, it's equivalent to prove the independence between $U_{\boldsymbol{\theta}_{\mathrm{f}}}(\hat{\boldsymbol{\lambda}})$ and $U_u(\hat{\boldsymbol{\tau}})$ by treating $\tilde{\mathbf{I}}_{\mathrm{f}}$ as fixed. Through Taylor expansion,

$$\mathbf{Y} - \mathbf{P}_{\mathrm{f}} \approx (\mathbf{I} - \mathbf{P}_1)(\mathbf{Y} - \mathbf{P}), \tag{14}$$

where $\mathbf{P}_1 = \mathbf{W}_{\mathrm{f}} \mathbf{X}_M \mathbf{Z}_{\mathbf{X}} (\mathbf{Z}_{\mathbf{X}}^T \mathbf{X}_M^T \mathbf{W}_{\mathrm{f}} \mathbf{Z}_{\mathbf{X}} \mathbf{X}_M)^{-1} \mathbf{Z}_{\mathbf{X}}^T \mathbf{X}_M^T$, $\mathbf{I}$ is the identity matrix. Similarly we have,

$$\mathbf{Y} - \mathbf{P}_{\mathrm{r}} \approx (\mathbf{I} - \mathbf{P}_2)(\mathbf{Y} - \mathbf{P}), \tag{15}$$

where $\mathbf{P}_2 = \mathbf{W}_{\mathrm{r}} \mathbf{C}_M \mathbf{Z}_{\boldsymbol{\tau}} (\mathbf{Z}_{\boldsymbol{\tau}}^T \mathbf{C}_M^T \mathbf{W}_{\mathrm{f}} \mathbf{Z}_{\boldsymbol{\tau}} \mathbf{C}_M)^{-1} \mathbf{Z}_{\boldsymbol{\tau}}^T \mathbf{C}_M^T$. Then we have $\mathbf{Z}_{\boldsymbol{\tau}}^T \mathbf{C}_M^T \mathbf{W}_{\mathrm{r}} (\mathbf{I} - \mathbf{P}_2)^T = \mathbf{0}$. Since $\mathbf{C} = [\mathbf{G}, \mathbf{X}]$ and $\mathbf{Z}_{\boldsymbol{\tau}} = \{\mathbf{Z}_{\mathrm{f}} \oplus (\oplus_{p=1}^{P} \mathbf{Z}^{(p)})\}_{\pi}$, then $\mathbf{C}_M \mathbf{Z}_{\boldsymbol{\tau}}$ can be written as $[\mathbf{G}_M \mathbf{Z}_{\mathrm{f}}, \mathbf{X}_M \mathbf{Z}_{\boldsymbol{\tau}}]$. Because $\mathbf{Z}_{\boldsymbol{\tau}}^T \mathbf{C}_M^T \mathbf{W}_{\mathrm{r}} (\mathbf{I} - \mathbf{P}_2)^T = \mathbf{0}$, then

$$\begin{aligned} \mathbf{Z}_{\mathrm{f}}^T \mathbf{G}_M^T \mathbf{W}_{\mathrm{r}} (\mathbf{I} - \mathbf{P}_2)^T &= \mathbf{0}, \\ \mathbf{Z}_{\mathbf{X}}^T \mathbf{X}_M^T \mathbf{W}_{\mathrm{r}} (\mathbf{I} - \mathbf{P}_2)^T &= \mathbf{0}. \end{aligned} \tag{16}$$

Through central limit theorem,

$$(U_{\boldsymbol{\theta}_{\mathrm{f}}}(\hat{\boldsymbol{\lambda}})^T, U_u(\hat{\boldsymbol{\tau}})^T)^T \xrightarrow{d} N(\mathbf{0}, \boldsymbol{\Sigma}), \tag{17}$$

where $\Sigma$ are the covariance matrix. To prove the independence between $U_{\boldsymbol{\theta}_{\mathrm{f}}}(\hat{\boldsymbol{\lambda}})$ and $U_u(\hat{\boldsymbol{\tau}})$, we only need to prove $cov\{U_{\boldsymbol{\theta}_{\mathrm{f}}}(\hat{\boldsymbol{\lambda}}), U_u(\hat{\boldsymbol{\tau}})\} = \mathbf{0}$. Hence

$$\begin{aligned}
cov\{U_{\boldsymbol{\theta}_{\mathrm{f}}}(\hat{\boldsymbol{\lambda}}), U_u(\hat{\boldsymbol{\tau}})\} &\approx cov\big\{\mathbf{Z}_{\mathrm{f}}^T\mathbf{G}_M^T(I-\mathbf{P}_1)(\mathbf{Y}-\mathbf{P}), \mathbf{Z}_{\mathrm{r}}^T\mathbf{G}_M^T(\mathbf{I}-\mathbf{P}_2)(\mathbf{Y}-\mathbf{P})\big\} \\
&= \mathbf{Z}_{\mathrm{f}}^T\mathbf{G}_M^T(\mathbf{I}-\mathbf{P}_1)\mathbf{W}_{\mathrm{r}}(\mathbf{I}-\mathbf{P}_2)^T\mathbf{G}_M\mathbf{Z}_{\mathrm{r}} \\
&= -\mathbf{P}_1\mathbf{W}_{\mathrm{r}}(\mathbf{I}-\mathbf{P}_2)^T\mathbf{G}_M\mathbf{Z}_{\mathrm{r}} \\
&= \mathbf{0}. \qquad (18)
\end{aligned}$$

### 5 Computation time simulations

We compared the computation time of analyzing 1,000 SNPs for five different methods: FTOP, MTOP, standard logistic regression, FTOP with only complete data and polytomous logistic regression. The methods were split into two groups: 1. Methods working on all the data consisted of MTOP, FTOP, standard logistic regression. 2. Methods working on complete data consisted of FTOP with complete data and polytomous model. To have a fair comparison with the polytomous model, FTOP with complete data was implemented using the Wald test so that the MLE estimation time is included. We generated the data based on the four and six tumor characteristics setting described in Section 3.2.1 in the main manuscript. The total sample size was set to be 5,000, 25,000, 50,000 and 100,000. The cases with complete data were respectively around 30% and 23% for the four and six tumor characteristics settings. We performed 1,000 independent simulations to calculate the averaged processing time for each method. All analyses were implemented in R version 3.6.0. MTOP, FTOP were implemented in TOP package version 1.0.8. Standard logistic regression was carried out in stats package version 3.6.1. (**?**). Polytomous model used nnet package 7.3.12 (**?**). The polytomous model in R was implemented in two steps. The first step used multinom function to fit the model. The second step used vcov function to get the covariance matrix of the estimates. Around 30% computation time was in the second step which could be potentially due to inefficient implementation of vcov function in R.

Supplementary Figure 1 presents the log of averaged computation times (s) for 1000 SNPs. Standard logistic regression had the smallest computation time among all the methods since standard logistic regression didn't include any tumor marker data in the analysis. FTOP was significantly faster compared to MTOP, since FTOP only needed to estimate parameters for all the covariates under the null hypothesis for one time. FTOP was 32-71 fold computationally faster than MTOP under different simulation settings. FTOP with only complete data was 2-14 fold computationally faster than the polytomous model under different simulation settings. More discussion about the computation complexity of MTOP and FTOP could be found in Section 5 of the main manuscript.

#### 6 Bias evaluation

We conducted simulations to evaluate whether removing the rare subtypes with less than 10 cases would bias the estimates. Similar setting as Section 3 in the main manuscript, we simulated the data mimicking the PBCS. Under the four tumor characteristics setting, the effect sizes were 0.08, 0.08, 0.05, 0.05, 0.05 for baseline effect, ER, PR, HER2, and grade main effect, respectively. Under the six tumor characteristics setting, two additional tumor characteristics were simulated with an effect as 0.05. For each simulation, we removed subtypes with less than 10 cases, just as in our previous simulations and real data analysis. We generated 10,000 simulation replicates to evaluate the bias of the two-stage model estimate after removing the rare subtypes.

As Supplementary Table 3 shows, the estimates were unbiased in both four and six tumor characteristics settings. Since removing the rare subtypes didn't involve the genetic effects, the estimates were unbiased.

#### 7 Power simulation with 5,000 subjects

Additional simulations for global association tests with 5,000 subjects were also considered for a larger effect size. We set $\beta_m$ as 0.25 for the scenario I, when there was no heterogeneity between tumor markers. The case-case parameter for ER ($\theta_1^{(1)}$) as 0.25 for scenario II, when heterogeneity was only driven by ER. The case-case parameter for ER ($\theta_1^{(1)}$) as 0.25, and the other case-case parameters were set to follow a normal distribution with mean 0 and variance $4.0\times10^{-4}$, when there were multiple tumor markers driving the heterogeneity. Similar to the low effect size simulations, standard logistic regression had the highest power when there presented no heterogeneity across subtypes.similar to the lower effect size simulations, MTOP had the highest power when there were heterogeneous associations (Supplementary Figure 2). The power of FTOP with only complete data and polytomous model were almost 0 under this case.

Table 1: The correlations of ER, PR, HER2, and grade

| Correlation | ER | PR | HER2 | Grade |
|---|---|---|---|---|
| ER | 1 | 0.61 | -0.16 | -0.39 |
| PR | 0.61 | 1 | -0.17 | -0.32 |
| HER2 | -0.16 | -0.17 | 1 | 0.20 |
| Grade | -0.39 | -0.32 | 0.20 | 1 |

Table 2: The frequencies of the joint distribution of ER (positive vs. negative), PR (positive vs. negative), HER2(positive vs. negative), and grade (ordinal 1, 2, 3). Within the table, "-" and "+" represent negative and positive, respectively.

| ER | PR | HER2 | Grade | Frequency (%) |
|---|---|---|---|---|
| - | - | - | 1 | 0.38 |
| + | - | - | 1 | 2.42 |
| - | + | - | 1 | 0.21 |
| + | + | - | 1 | 15.04 |
| - | - | + | 1 | 0.09 |
| + | - | + | 1 | 0.24 |
| - | + | + | 1 | 0.02 |
| + | + | + | 1 | 0.87 |
| - | - | - | 2 | 2.64 |
| + | - | - | 2 | 5.87 |
| - | + | - | 2 | 0.71 |
| + | + | - | 2 | 32.74 |
| - | - | + | 2 | 1.31 |
| + | - | + | 2 | 1.37 |
| - | + | + | 2 | 0.16 |
| + | + | + | 2 | 3.89 |
| - | - | - | 3 | 9.50 |
| + | - | - | 3 | 3.02 |
| - | + | - | 3 | 0.69 |
| + | + | - | 3 | 10.30 |
| - | - | + | 3 | 3.75 |
| + | - | + | 3 | 1.39 |
| - | + | + | 3 | 0.28 |
| + | + | + | 3 | 3.11 |

Table 3: The bias of the estimates from the two-stage polytomous model using the EM algorithm with $10^5$ randomly simulated samples. T5 and T6 represent the two additional binary tumor characteristics simulated other than ER, PR, HER2 and grade in the six tumor characteristics setting.

| Sample Size | ER | PR | HER2 | Grade | T5 | T6 |
|---|---|---|---|---|---|---|
| | | | Four tumor characteristics | | | |
| 25,000 | $-2 \times 10^{-3}$ | $1 \times 10^{-3}$ | $1 \times 10^{-3}$ | 0.00 | | |
| 50,000 | 0.00 | 0.00 | 0.00 | 0.00 | | |
| 100,000 | 0.00 | 0.00 | 0.00 | 0.00 | | |
| | | | Six tumor characteristics | | | |
| 25,000 | 0.00 | $-2 \times 10^{-3}$ | $-4 \times 10^{-3}$ | $-1 \times 10^{-3}$ | 0.00 | 0.00 |
| 50,000 | 0.00 | 0.00 | 0.00 | 0.00 | 0.00 | 0.00 |
| 100,000 | 0.00 | 0.00 | 0.00 | 0.00 | 0.00 | 0.00 |

Table 4: Sample size of four tumor characteristics in Polish Breast Cancer Study

| | ER | PR | HER2 | | Grade |
|---|---|---|---|---|---|
| Positive | 1,316 | 1,056 | 1,246 | Grade 1 | 356 |
| Negative | 594 | 847 | 254 | Grade 2 | 968 |
| Missing | 168 | 157 | 578 | Grade 3 | 554 |
| | | | | Missing | 200 |

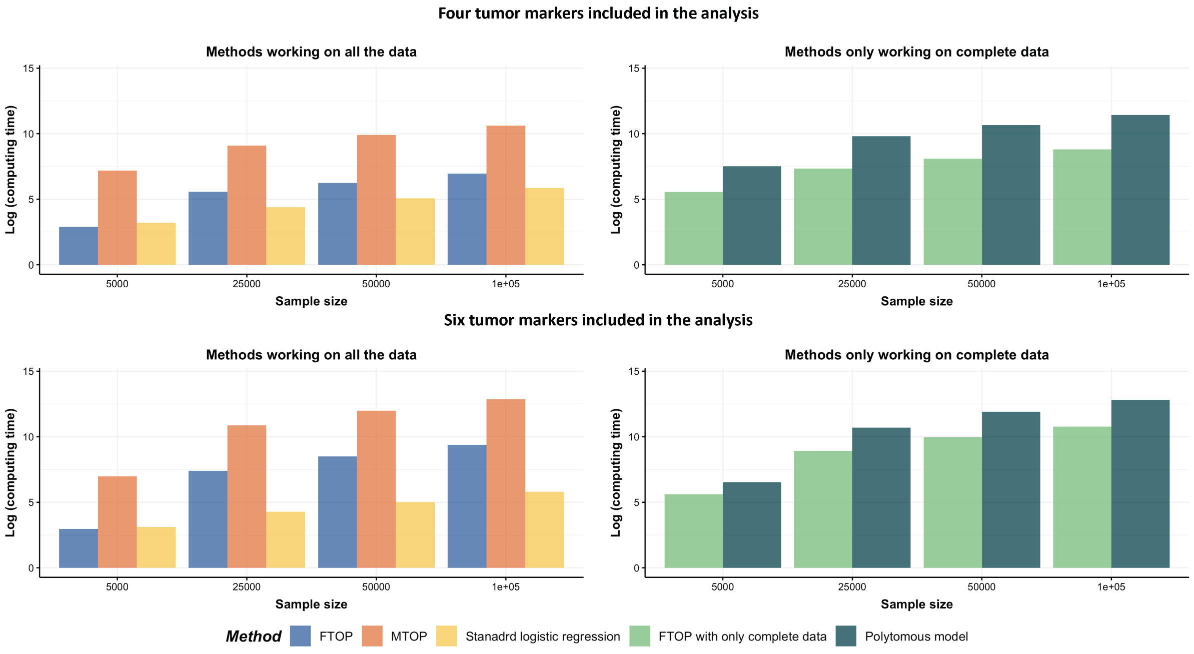


Figure 1: The log of computing time(s) for 1,000 SNPs. The computation time for MTOP, FTOP, standard logistic regression, the two-stage model with only complete data and polytomous model were estimated using $1,000$ random samples. In the first row, four tumor markers were included in the analysis. Three binary tumor markers and one ordinal tumor marker defined 24 cancer subtypes. In the second row, two extra binary tumor markers were included in the analysis. The six tumor markers defined 96 subtypes. Around 77% cases would be incomplete.

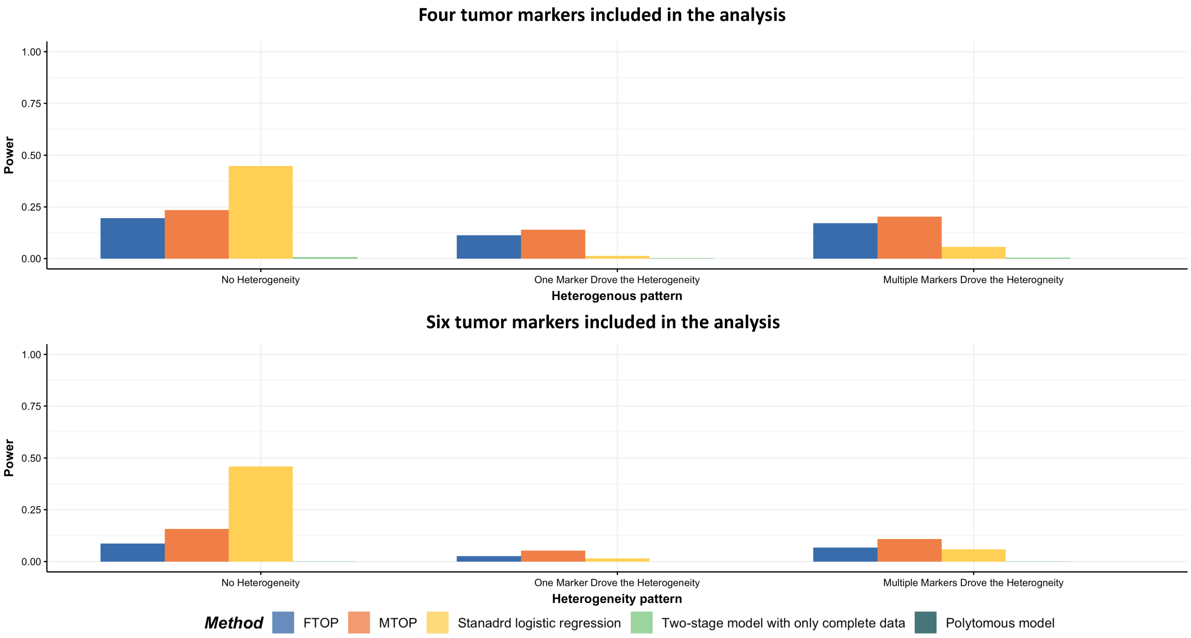


Figure 2: Global association test power simulations using MTOP, FTOP, standard logistic regression, FTOP with only complete data and polytomous model with 5,000 subjects. For the figure in the first row, four tumor markers were included in the analysis. Three binary tumor markers and one ordinal tumor marker defined 24 cancer subtypes. Around 70% cases were incomplete. For the three figures in the second row, two extra binary tumor markers were included in the analysis. The six tumor markers defined 96 subtypes. Around 77% cases were incomplete. We generated $2 \times 10^5$ random simulation replicates. The power was estimated by controlling the type I error $\alpha < 5.0 \times 10^{-8}$. The power for FTOP with complete data and polytomous model were almost 0.

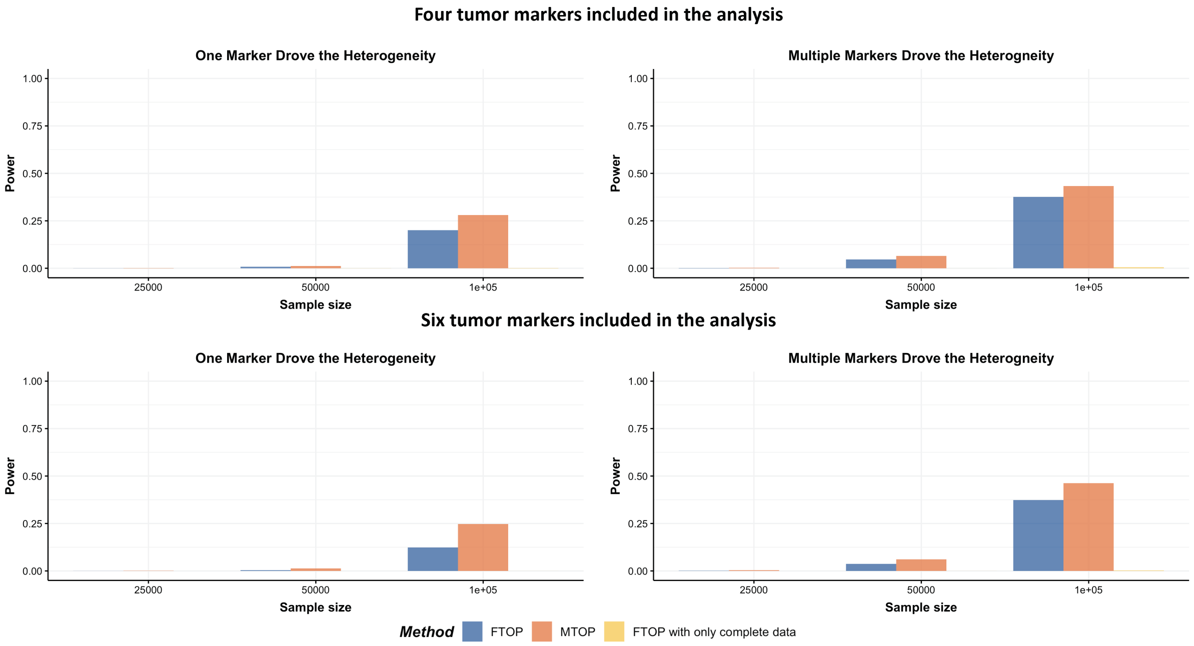


Figure 3: Global heterogeneity test power simulations result using FTOP, MTOP, and FTOP with only complete data. For the two figures in the first row, four tumor markers were included in the analysis. Three binary tumor markers and one ordinal tumor marker defined 24 cancer subtypes. Around 70% cases would be incomplete. For the two figures in the second row, two extra binary tumor markers were included in the analysis. The six tumor markers defined 96 subtypes. Around 77% cases would be incomplete. The total sample size was 25,000, 50,000 and 100,000. We generated $2 \times 10^5$ random simulation replicates. The power was estimated by controlling the type I error $\alpha < 5.0 \times 10^{-8}$. The power for FTOP with complete data was almost 0.

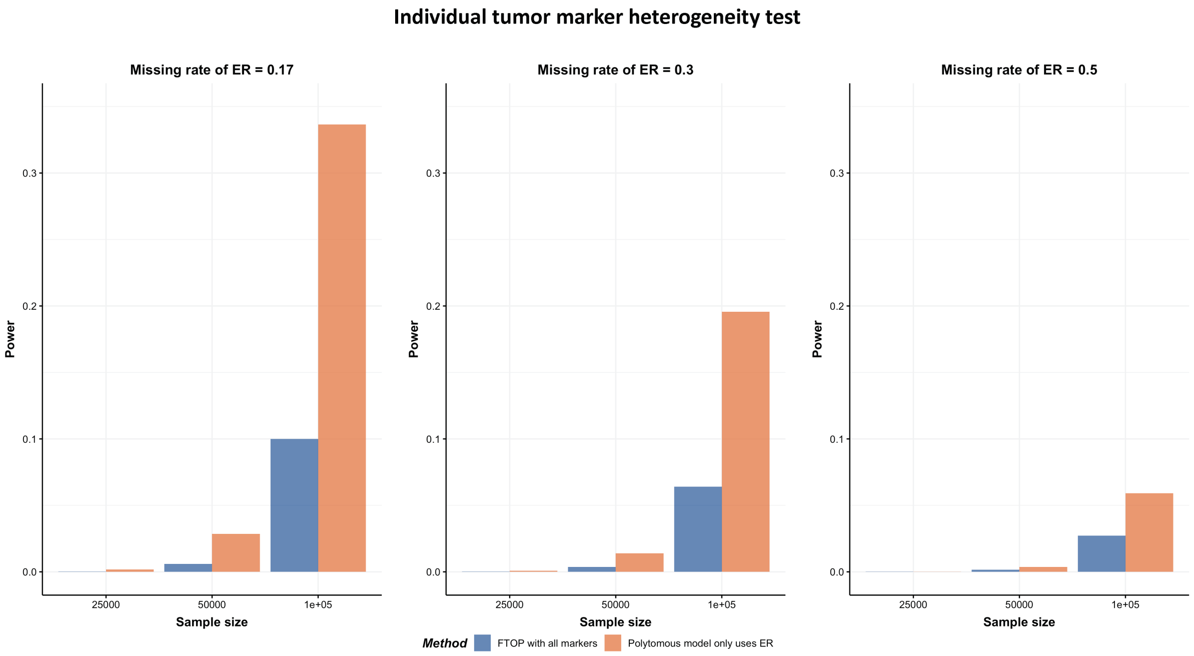


Figure 4: Individual tumor marker heterogeneity test power simulation results using FTOP with all the markers, polytomous model with only ER. Four tumor markers were included to generate the datasets. Three binary tumor markers and one ordinal tumor marker defined 24 cancer subtypes. The missing rate for ER was 0.17, 0.30 and 0.50. The effect of ER was 0.08, and the effects of PR, HER2 and grade were 0. The total sample size was 25,000, 50,000 and 100,000. We generated $2 \times 10^5$ independent simulations replicates. The power was estimated by controlling the type I error $\alpha < 5.0 \times 10^{-8}$.

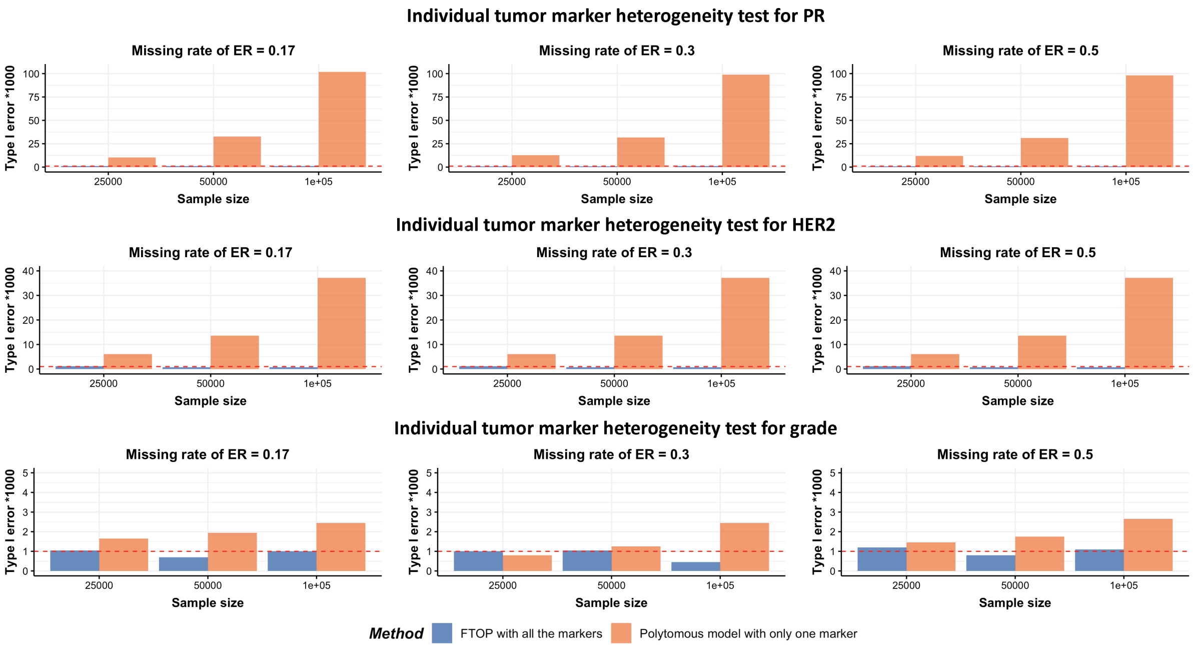


Figure 5: Type I error of individual tumor marker heterogeneity test for PR, HER2, and grade using FTOP with all the four markers, the polytomous model with only one marker at a time. Four tumor markers were included to generate the datasets. Three binary tumor marker and one ordinal tumor marker defined 24 cancer subtypes. The missing rate for ER was 0.17, 0.30 and 0.50. The effect of ER was 0.08, and the effects of PR, HER2 and grade were 0. The total sample size was 25,000, 50,000 and 100,000. We generated $2 \times 10^5$ independent simulations replicates. The type I error was evaluated at $1.0 \times 10^{-3}$ level given the number of simulation replicates. The red dashed line showed the corrected type I error.

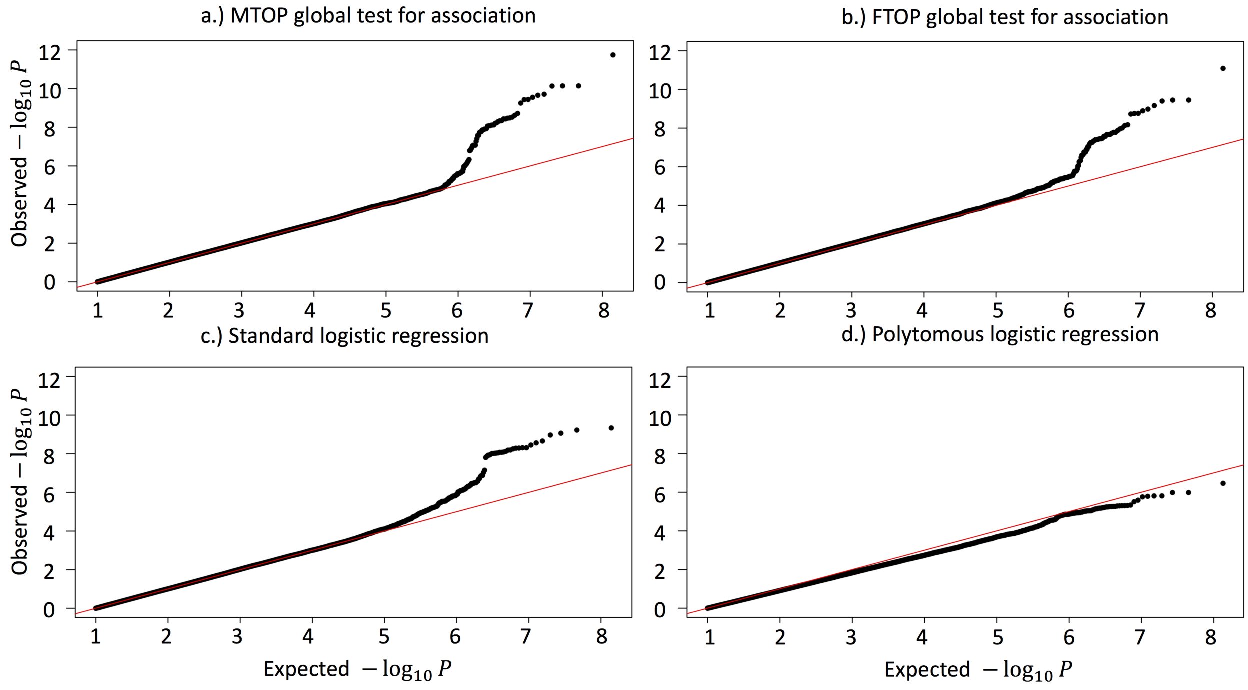


Figure 6: QQ plot of genome-wide association analysis with PBCS using MTOP, FTOP, standard logistic regression, the polytomous model. MTOP and FTOP used additive structure. MTOP assumed baseline and ER effects as fixed effects, and all the other effects were assumed as random effects. PBCS had 2,078 invasive breast cancer and 2,219 controls. In total, 7,017,694 SNPs on 22 auto chromosomes with MAF more than 5% were included in the analysis. ER, PR, HER2, and grade were used to define breast cancer subtypes.

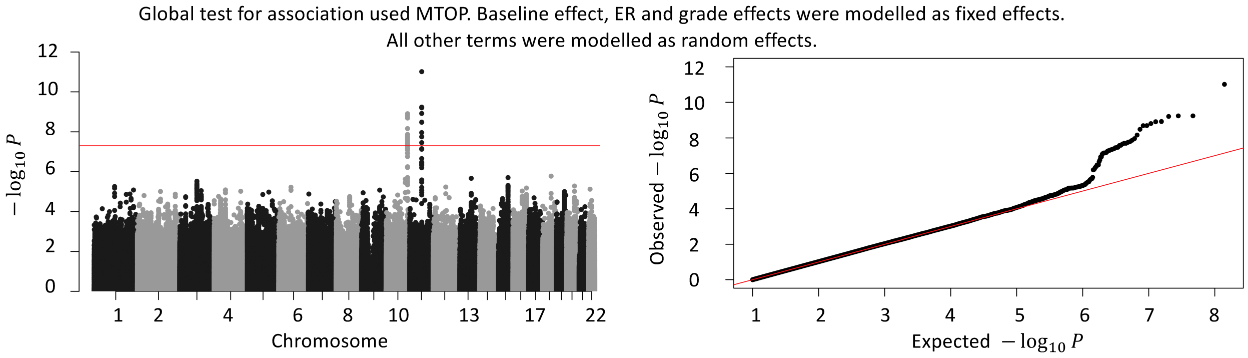


Figure 7: Manhattan plot and QQ plot of genome-wide association analysis with PBCS using MTOP with additive structure. The baseline effect, ER and grade effects were modeled as fixed effects. PBCS had 2,078 invasive breast cancer cases and 2,219 controls. In total, 7,017,694 SNPs on 22 auto chromosomes with MAF more than 5% were included in the analysis. ER, PR, HER2, and grade were used to define breast cancer subtypes.

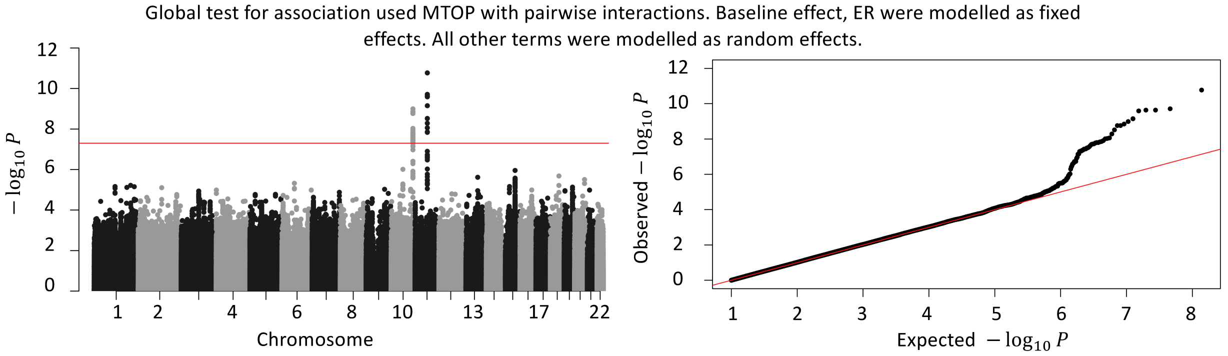


Figure 8: Manhattan plot and QQ plot of genome-wide association analysis with PBCS using MTOP with pairwise interactions. Baseline effect and ER effect were modeled as fixed effects. All the other effects were modeled as random effects. PBCS had 2,078 invasive breast cancer cases and 2,219 controls. In total, 7,017,694 SNPs on 22 auto chromosomes with MAF more than 5% were included in the analysis. ER, PR, HER2, and grade were used to define breast cancer subtypes.

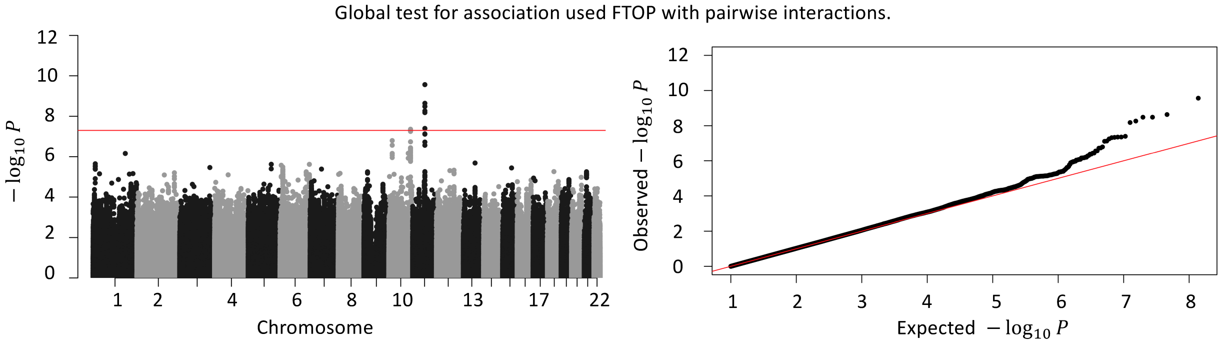


Figure 9: Manhattan plot and QQ plot of genome-wide association analysis with PBCS using FTOP with pairwise interactions. PBCS has 2,078 invasive breast cancer cases and 2,219 controls. In total, 7,017,694 SNPs on 22 auto chromosomes with MAF more than 5% were included in the analysis. ER, PR, HER2, and grade were used to define breast cancer subtypes.
